# Malaria transmission relies on concavin-mediated maintenance of *Plasmodium* sporozoite cell shape

**DOI:** 10.1101/2021.11.06.467543

**Authors:** Jessica Kehrer, Pauline Formaglio, Julianne Mendi Muthinja, Sebastian Weber, Danny Baltissen, Christopher Lance, Johanna Ripp, Janessa Grech, Markus Meissner, Charlotta Funaya, Rogerio Amino, Friedrich Frischknecht

## Abstract

During transmission of malaria-causing parasites from mosquitoes to mammals, *Plasmodium* sporozoites migrate rapidly in the skin to search for a blood vessel. The high migratory speed and narrow passages taken by the parasites suggest considerable strain on the sporozoites to maintain their shape. Here we report on a newly identified protein, concavin, that is important for maintenance of the sporozoite shape inside salivary glands of mosquitoes and during migration in the skin. Concavin-GFP localized at the cytoplasmic periphery of sporozoites and *concavin(−)* sporozoites progressively rounded up upon entry of salivary glands. These rounded *concavin(−)* sporozoites failed to pass through the narrow salivary ducts and were hence rarely ejected by mosquitoes. However, normally shaped *concavin(−)* sporozoites could be transmitted and migrated in the skin or skin like environments. Strikingly, motile *concavin(−)* sporozoites could disintegrate while migrating through narrow strictures in the skin leading to parasite arrest or death and decreased transmission efficiency. We suggest that concavin contributes to cell shape maintenance by riveting the plasma membrane to the subtending inner membrane complex.

**SIGNIFICANCE:** Malaria parasites are transmitted by *Anopheles* mosquitoes and rely on rapid migration for establishing an infection. We identified and characterized a protein, named concavin, essential for maintaining the shape of the sporozoite. Concavin is a membrane associated protein facing the cytoplasm suggesting that it contributes to riveting the plasma membrane to the subtending inner membrane complex. Sporozoites lacking concavin can round up in the salivary glands, are less well transmitted to mice and disintegrate while migrating in the skin. Hence, concavin is essential for parasite transmission and infectivity.

**Highlights:**

- A membrane associated protein is essential for *Plasmodium* shape maintenance
-Migrating parasites disintegrate in the absence of concavin
-First protein essential for cellular integrity of *Plasmodium* sporozoites
-Thickened and deformed *Plasmodium* sporozoites fail to be transmitted by mosquitoes

## INTRODUCTION

Malaria is still prevalent in tropical countries where it infects over 200 million people every year killing over 400,000, mostly young African children (WHO, 2020). While the symptoms of the disease are caused by the parasite stages infecting red blood cells, the only licenced malaria vaccine has been derived from the surface circumsporozoite protein (CSP) of the mosquito-transmitted parasite stage, the *Plasmodium* sporozoite (Clemens & Moorthy, 2016; Cowman et al., 2016). CSP is essential for sporozoite formation and functions at different steps of the sporozoite journey from the mosquito gut to the mammalian liver (Aliprandini et al., 2018; Coppi et al., 2011; Ménard et al., 1997; Singer & Frischknecht, 2021; Thathy et al., 2002). Unfortunately, the CSP-based vaccine has failed to deliver the long-sought efficient protection from malaria infections and new vaccine candidates are urgently needed for exploration (Matuschewski, 2017). Secreted and/or plasma membrane associated sporozoite proteins might constitute additional good candidates for next generation vaccines.

*Plasmodium* sporozoites are deposited into the dermis during a mosquito bite and migrate at high speed to enter both blood or lymph vessels (Amino et al., 2006; Hopp et al., 2021). Those entering the blood can ultimately exit the circulation in the liver and infect hepatocytes to further develop into red blood cell infecting merozoites (Prudêncio et al., 2006; Tavares et al., 2013). Sporozoites are highly polarized and slender cells with a chiral sub-pellicular cytoskeleton that defines parasite length and curvature linked to the inner membrane complex (IMC) that subtends the plasma membrane (Gould et al., 2008; Harding & Frischknecht, 2020; Khater et al., 2004; Kudryashev et al., 2012; Spreng et al., 2019; Tremp et al., 2013; Volkmann et al., 2012). The IMC is found at a constant distance to and hence likely linked to the plasma membrane as shown for *Toxoplasma gondii* (Frénal et al., 2010). Disruption of some IMC-proteins leads to parasite swelling around the nucleus, which impacts motility and infectivity (Khater et al., 2004; Volkmann et al., 2012). However, many proteins of the pellicle remain to be described and no detailed picture is yet available about how these complex interactions form and maintain the cellular shape (Harding & Frischknecht, 2020).

Sporozoites secrete proteins from micronemal vesicles and rhoptries at their apical pole (Dubremetz et al., 1998). Like CSP, these proteins can be essential for the escape of sporozoites from oocysts at the *Anopheles* midgut wall into the mosquito hemolymph, to enter salivary glands, for migration within the skin, to enter blood vessels and ultimately hepatocytes (Carey et al., 2014; Ishino et al., 2019; Klug & Frischknecht, 2017; Risco-Castillo et al., 2015; Silvie et al., 2004; Wang et al., 2005). Sporozoite migration within the skin provides the first possible target for intervention against an infection with *Plasmodium* (Aliprandini et al., 2018; Douglas et al., 2018; Murugan et al., 2020). Antibodies targeting CSP can inhibit sporozoite motility (Aliprandini et al., 2018; Vanderberg & Frevert, 2004) and induce self-killing of sporozoites via secreted pore-forming proteins, which are essential for sporozoite migration through cells (Aliprandini et al., 2018; Amino et al., 2008; Bhanot et al., 2003; Risco-Castillo et al., 2015). Other secreted proteins include TRP1, LIMP and CelTOS as well as members of the transmembrane TRAP (thrombospondin related anonymous protein) family. TRP1 (thrombospondin related protein 1) is essential for the release of sporozoites from oocysts and invasion of salivary glands (Klug & Frischknecht, 2017). LIMP and TRAP are essential for sporozoite invasion of salivary glands and liver cells (Santos et al., 2017; Sultan et al., 1997) and CelTOS (cell traversal protein for ookinetes and sporozoites) is important for sporozoite motility and migration through cells (Jimah et al., 2016; Kariu et al., 2006; Steel et al., 2018). CelTOS and TRAP are investigated as possible vaccine candidates (Pirahmadi et al., 2019; Tiono et al., 2018).

Proteomic analysis of sporozoites revealed a number of yet uncharacterized proteins including proteins on the parasite surface (Lindner et al., 2019; Lindner, Swearingen, et al., 2013; Swearingen et al., 2016). However, identification of surface proteins has been a challenge due to contamination from cytoplasmic proteins. We previously collected the supernatant from sporozoites activated by the addition of pluronic acid, which stimulates secretion (Kehrer, Singer, et al., 2016) and determined the proteins by mass spectrometry. From this set of putatively secreted proteins we selected the uncharacterized PbANKA_1422900 protein for further analysis by gene deletion and GFP-tagging and found that this protein is localized at the cytosolic side of the plasma membrane. PbANKA_1422900 is important for the maintenance of the sporozoite shape during their salivary gland residency and essential for efficient transmission to the vertebrate host. While deformed sporozoites could still move, they were not ejected through the narrow salivary ducts and failed to penetrate through skin and skin-like environments showing the importance of the slender shape for parasite transmission. Due to its impact on the convex-concave polarity of sporozoites we named the protein concavin. Strikingly, during sporozoite migration through strictures in the skin, we observed *concavin(−)* parasites to disintegrate by the apparent shedding of large membrane-delimited parts of the parasite.

## RESULTS

### Concavin is a conserved Apicomplexan protein important for sporozoite shape maintenance

PBANKA_1422900 is expressed highest in ookinetes and sporozoites according to RNA seq data obtained from Plasmodb.org (Figure S1). It is a 393 amino acid long protein in *P. berghei* and conserved among Apicomplexa. Concavin shares 96% amino acid residue identity with *P. yoelii*, 80% identity with *P. vivax*, 76% with *P. falciparum*, and 36% identity with the orthologue from *T. gondii* (Figure S1–S2). The only recognizable feature in this protein was a potential palmitoylation site at the N-terminus (Figure S2A). To test for a function of concavin, we disrupted the gene through double homologous recombination in *P. berghei* (Figure S3) and *T. gondii* (Figure S4). Deletion of concavin readily yielded clonal *P. berghei* parasite lines, which grew at similar multiplication rates per 24h as wild type parasites in the blood stage (Figure 1A). Similarly, we could not detect a phenotypic difference in a plaque assay between wild type and transgenic *T. gondii* (Figure S4). Transmission of the *P. berghei concavin(−)* parasites to mosquitoes showed slightly reduced numbers of oocysts in infected mosquitoes (Figure 1B). We regularly found large numbers of *concavin(−)* sporozoites within the salivary glands, however, a large proportion of sporozoites showed an abnormal shape. While wild type sporozoites usually keep the typical curved and slender shape at any time post salivary gland entry, *concavin(−)* sporozoites rounded up over time. Rounding up of *concavin(−)* sporozoites was initiated at the posterior end of the cell (Figure 1C) and hence appeared different to the rounding observed after liver cell entry (Jayabalasingham et al., 2010) (see also Figure 2E). This loss of curvature led us to name PbANKA_1422900 concavin. Curiously, over 90% of oocyst-derived *concavin(−)* sporozoites were normally formed. Yet with prolonged residency in salivary glands more sporozoites became deformed or rounded up completely (Figure 1D-E). In contrast, we never observed deformed wild type sporozoites, neither in the midgut nor in the salivary gland. Curiously, both deformed and normally shaped sporozoites were still able to move (Figure 1F-G). While normally shaped *concavin(−)* sporozoites displayed circular movement in a wild type manner with nearly wild type speed, deformed sporozoites progressed with significantly slower speed. (Figure 1F). Deformed sporozoites also moved on less curved paths as did wild type or normally formed *concavin(−)* parasites (Figure 1G).

**Figure 1.**
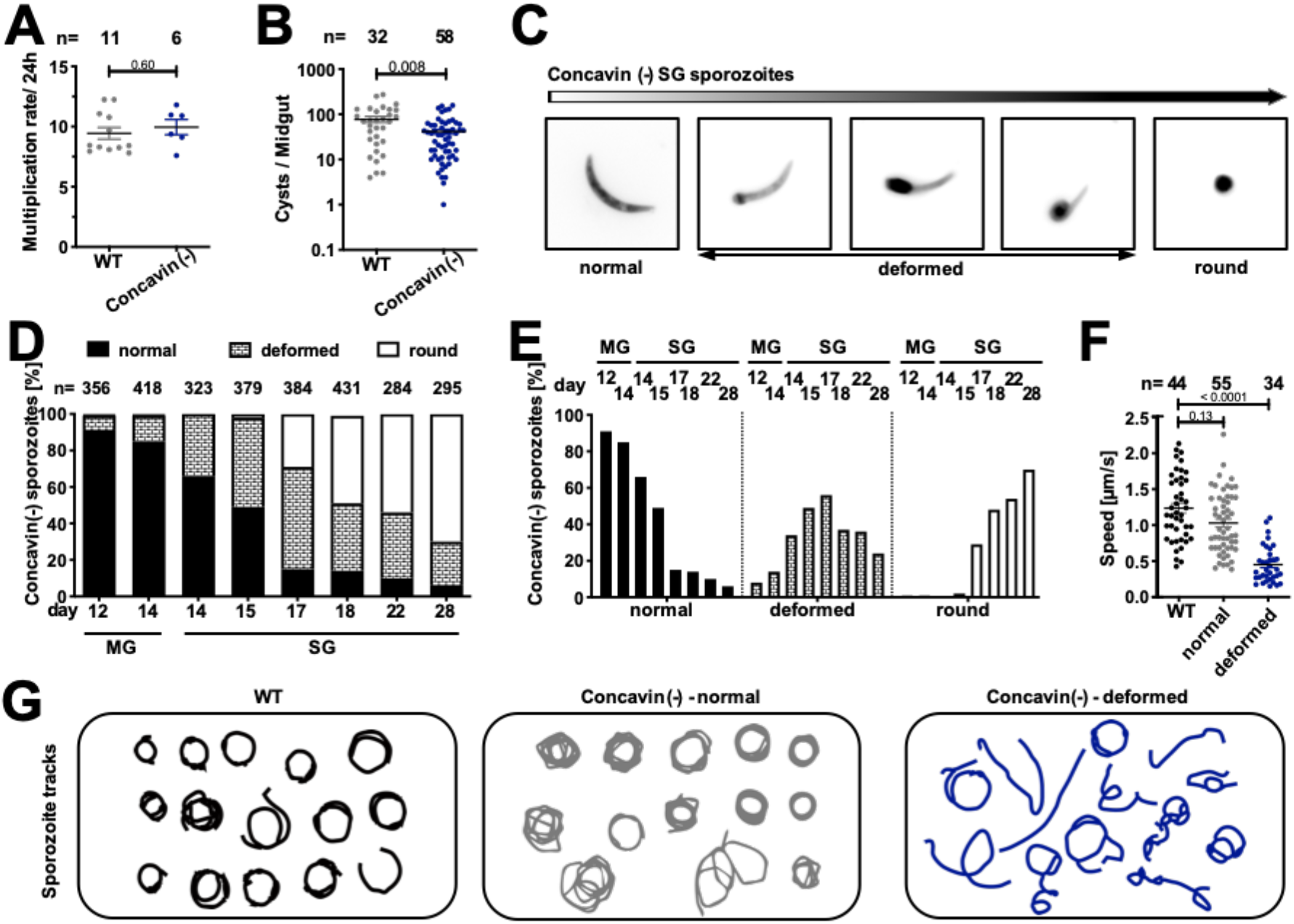
Deletion of *concavin* leads to rounding of sporozoites. **(A)** Blood-stage growth rate of *concavin(−)* parasites in comparison to wild type. Data points represent parasites growing in individual mice, n indicates total number. P-value is calculated using the Mann Whitney test. Shown is the mean ± SEM. **(B)** Oocyst development in the mosquito observed between d12-17 post infection. Data points represent individual midguts, n indicates total number. P-value is calculated using the Mann Whitney test. Shown is the mean ± SEM. **(C)** Mosquito infections resulted in deformed sporozoites in the salivary gland. Shown are example images of sporozoites arranged to illustrate their rounding up over time. **(D-E)** Quantification of sporozoite rounding up over time in the midgut (MG) and salivary gland (SG) at the indicated days post infection. Sporozoites were classified as either normal, deformed or round as illustrated in C. **(F)** Average speed of salivary gland sporozoites. Data points represent individual sporozoites, n indicates total number. P-values are calculated using the Kruskal Wallis test followed by the Dunns multiple comparison test. (G) Selected trajectories of manually tracked sporozoites.

**Figure 2.**
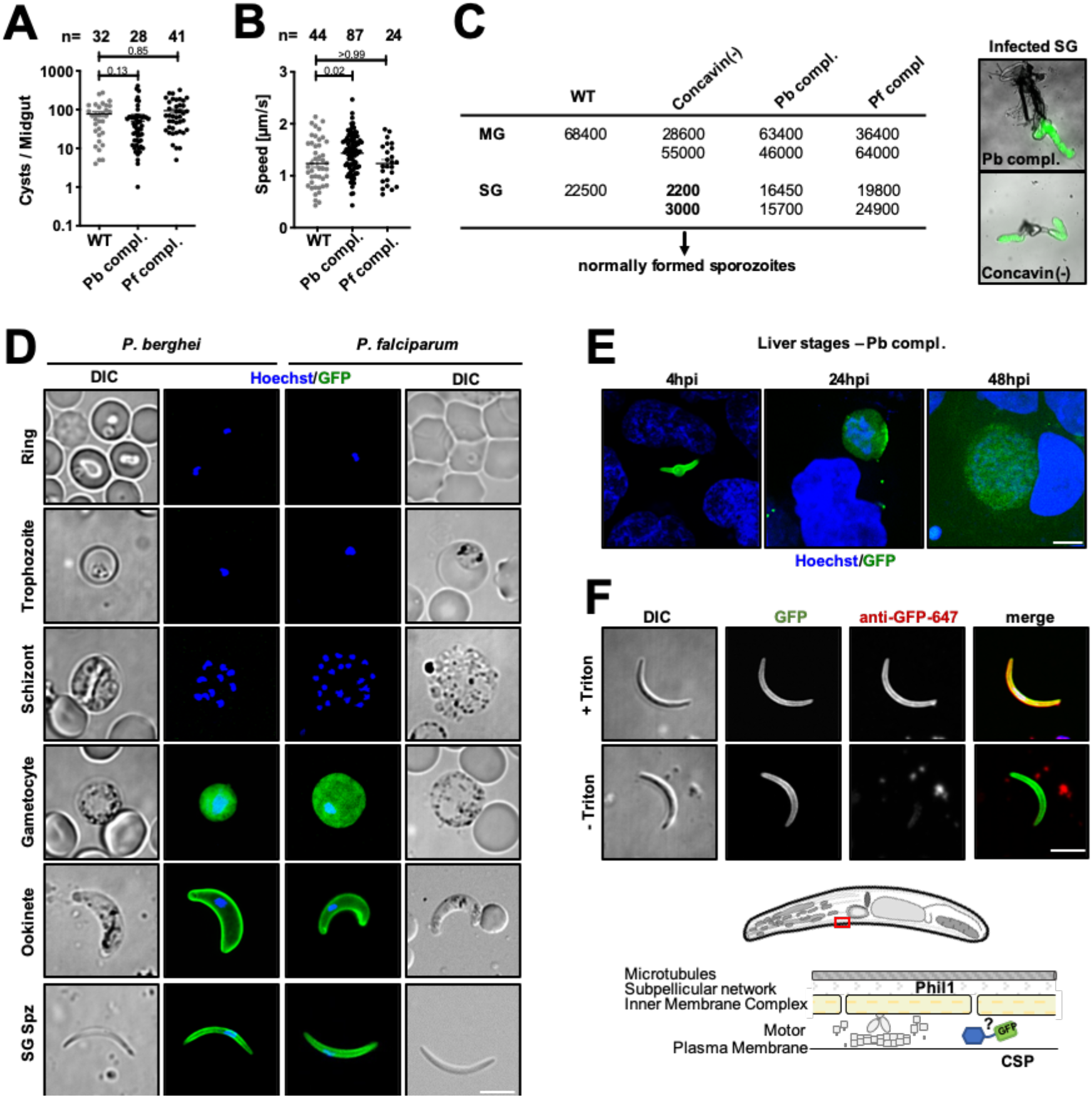
Concavin-GFP localizes to the periphery of ookinetes and sporozoites. **(A)** Oocyst development of *concavin(−)* parasites complemented with either the *P. berghei* gene or the *P. falciparum* orthologue fused to GFP. Data points represent individual midguts observed between d12-17 post infection. Shown is the mean ± SEM. P-values are calculated using the Kruskal Wallis test followed by the Dunns multiple comparison test. **(B)** Average speed of salivary gland sporozoites. Data points represent individual sporozoites. Shown is the mean ± SEM. P-values are calculated using the Kruskal Wallis test followed by the Dunns multiple comparison test. **(C)** Sum table of mosquito infections of wild type, *concavin(−)* and complemented lines expressing either *P. berghei* (Pb) or *P. falciparum* (Pf) concavin-GFP. Numbers determined on d17 post mosquito infection. For each counting at least 20 mosquitos were dissected. Please note for *concavin(−)* parasites only normally shaped sporozoites were counted. **(D)** Pb concavin-GFP and Pf concavin-GFP localization in blood and mosquito stages. Nuclei (blue) stained with Hoechst. Scale bar: 5 μm. **(E)** Localization of *P. berghei* concaving-GFP in liver stages. Nuclei (blue) stained with Hoechst. Scale bar: 5 μm. **(F)** Immunofluorescence images of permeabilized and unpermeabilized concavin-GFP expressing salivary gland sporozoites stained with an anti-GFP antibody. Note that a GFP signal could only be detected after permeabilization, excluding concavin localization on the parasite surface as illustrated in the model. Scale bar: 5μm.

### Concavin localizes at the cytosolic side of the periphery

To localize concavin, we generated two parasite lines expressing a GFP-tagged version of the protein. To this end, we first reintroduced the *P. berghei concavin-gfp* sequence into the knockout, thus complementing the *concavin(−)* parasite line. In addition, we also complemented the *concavin(−)* with the *P. falciparum concavin-gfp* gene (Figure S5) and we tagged the ortholog from *Toxoplasma gondii* in that parasite (Figure S4). Both *Plasmodium* lines were able to establish mosquito infections comparable to wild type levels including the colonization of salivary glands by highly motile sporozoites suggesting that both proteins are fully functional (Figure 2A-C). Concavin-GFP could be detected in gametocytes, ookinetes, sporozoites and liver stages and was absent in blood stage parasites (Figure 2D-E, supplementary movie 1). In gametocytes the protein localized diffusely while in ookinetes and salivary gland derived sporozoites concavin-GFP localized at the periphery suggesting an association with the plasma membrane. In *T. gondii* tachyzoites, we also found a peripheral signal (Figure S4). A peripheral localization could also derive from a protein resident in the sub-pellicular network, IMC or supra-alveolar space, the narrow space between IMC and the plasma membrane (Bane et al., 2016; Khater et al., 2004). To specify concavin-GFP localization we next fixed concavin-GFP expressing sporozoites and labelled them with anti-GFP antibodies with or without membrane permeabilization. Antibodies only detected concavin-GFP after permeabilization suggesting an internal localization of the protein (Figure 2F).

### Concavin-GFP is highly mobile and does not localize to the cytoskeleton

To test if concavin-GFP is associated with a cytoskeletal structure we employed fluorescence bleaching and monitored the recovery of the signal (FRAP). As a control we generated a PHIL1-GFP line (Figure S6) as PHIL1 is a constituent of the subpellicular network. This line recapitulated the published localization of the protein at the periphery of merozoites, ookinetes and sporozoites (Figure S6, (Saini et al., 2017)). Concavin-GFP expressing sporozoites showed a rapid recovery of the fluorescence signal after a bleached spot was introduced by a high energy laser suggesting high lateral diffusion of concavin-GFP (Figure 3A, supplementary movie 2). In contrast, in bleached PHIL1-GFP sporozoites there was no detectable recovery as expected from a protein anchored in the sub-pellicular network (Figure 3B, supplementary movie 2). These data suggest that concavin-GFP is not associated to the subpellicular network or another stable cytoskeletal structure.

**Figure 3.**
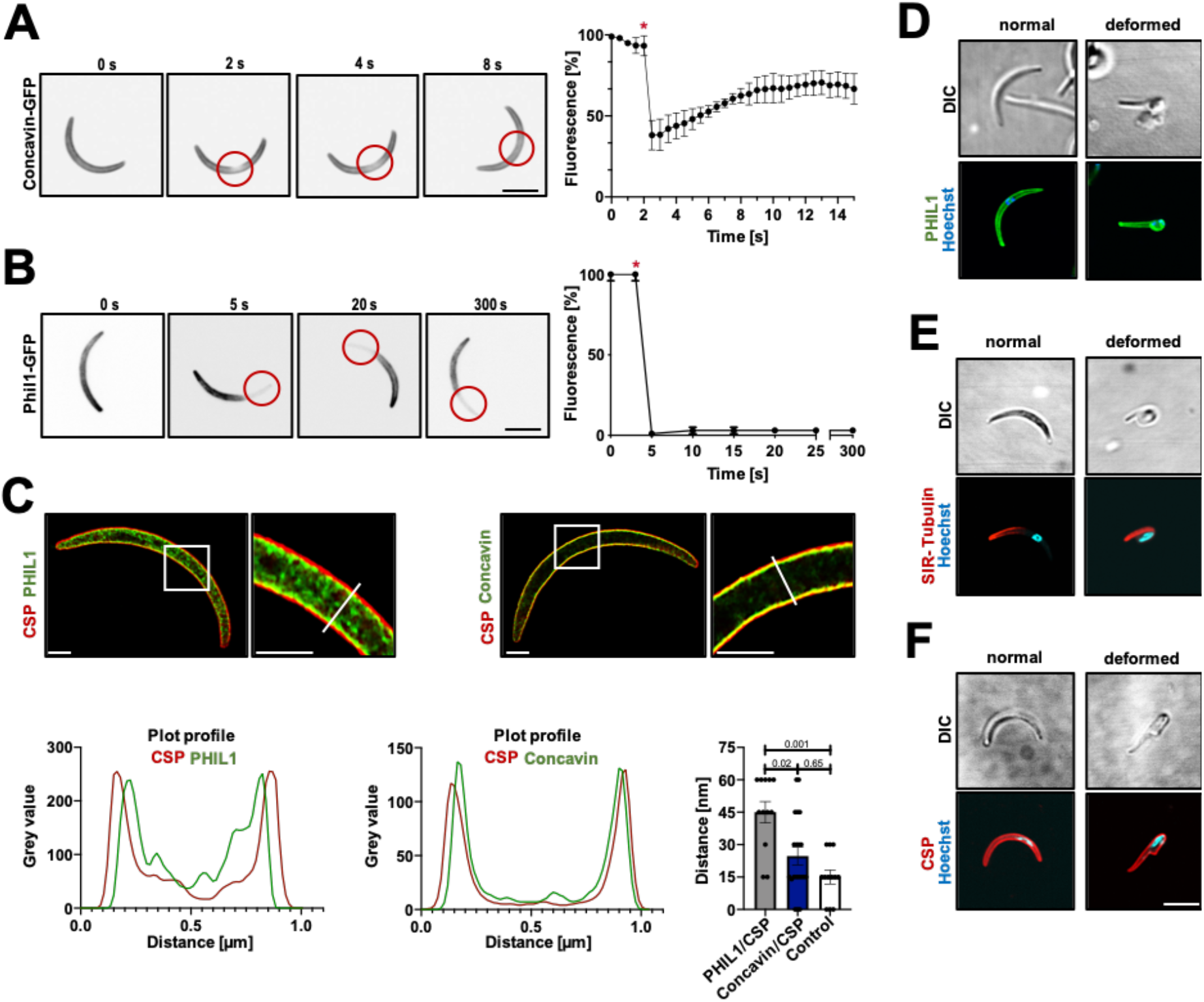
FRAP reveals dynamics of concavin-GFP and fixed PhiLl1-GFP. **(A, B**) Time series of concavin-GFP (A) PhiL1-GFP (B) sporozoites before and after bleaching and quantification of the fluorescence signal over time. * indicates time of bleaching. **(C)** Super resolution (STED) imaging of PhiL1-GFP and CSP as well as concavin-GFP and CSP. Cells were stained with an anti-GFP antibody in combination with Atto-594 (green) in addition to an anti-CSP staining in combination with Atto-647. Images were deconvolved using the Richardson-Lucy algorithm. The distance between the 2 signal peaks was measured using the plot profile of the respective channels in Fiji. Data points represent distance in individual sporozoites at the center of the cell. P-values are calculated using the Kruskal Wallis test followed by the Dunns multiple comparison test. Scale bar: 1 μm. **(D)** Localization of PhiL1-GFP (green) in *concavin(−)* parasites. Nuclei (blue) stained with Hoechst. **(E)** Localization of SiR-Tubulin (red) in *concavin(−)* parasites. Nuclei (blue) stained with Hoechst. **(F)** Localization of CSP (red) in *concavin(−)* parasites. Nuclei (blue) stained with Hoechst. Scale bar: 5 μm.

Next, we performed super-resolution (STED) co-localization experiments with antibodies against concavin-GFP, CSP and PHIL1-GFP. STED imaging showed that the signals of anti-GFP antibodies detecting PHIL1-GFP and antibodies against CSP were spatially separated (Figure 3C). In contrast, anti-GFP antibodies detecting concavin-GFP co-localized with anti-CSP antibodies. Similarly, antibodies recognizing CSP but stained with two different colours also co-localized (Figure 3C, Figure S7). This suggests that concavin-GFP is localized closer to the plasma membrane than PHIL1, probably within the alveolar space between IMC and plasma membrane or at the inner leaflet of the plasma membrane.

We next generated a non-clonal parasite line via single homologous recombination that expresses PHIL1-GFP in the *concavin(−)* parasite to investigate the subpellicular network in these parasites (Figure S6). PHIL1-GFP localization of this parasite was similar to that of PHIL1-GFP in wild type parasites in blood stages (Figure S6). However, in sporozoites, PHIL1-GFP appeared in an aberrantly localized manner, as would be expected from the changed shape of the sporozoite (Figure 3D, Figure S8). We compared this staining to *concavin(−)* parasites where the microtubules were labelled with SiR tubulin or with anti-CSP antibodies. This showed that microtubules appeared largely intact (Figure 3E, Figure S8) and that CSP was found on the surface of the deformed sporozoites (Figure 3F, Figure S8). Curiously, however, close inspection of the PHIL1-GFP labelling showed unexpected accumulations of signal. To investigate the deformed sporozoites at higher resolution, sporozoite containing salivary glands were examined by transmission and scanning electron microscopy. This showed the IMC subtending the plasma membrane in both wild type (Figure 4A) and *concavin(−)* sporozoites (Figure 4B-C). In rounded *concavin(−)* sporozoites, the IMC was in addition partially not associated with the plasma membrane anymore, but extended deep into the sporozoite cytoplasm (Figure 4B-C), Figure S9). Using array tomography we next reconstructed serial sections of a complete rounded *concavin(−)* sporozoite. This clearly revealed an intact tube-like ‘rolled up’ structure separated by the IMC from the remaining plasma membrane (Figure 4C). This suggests that the IMC is still intact but detached from the subtending plasma membrane.

**Figure 4.**
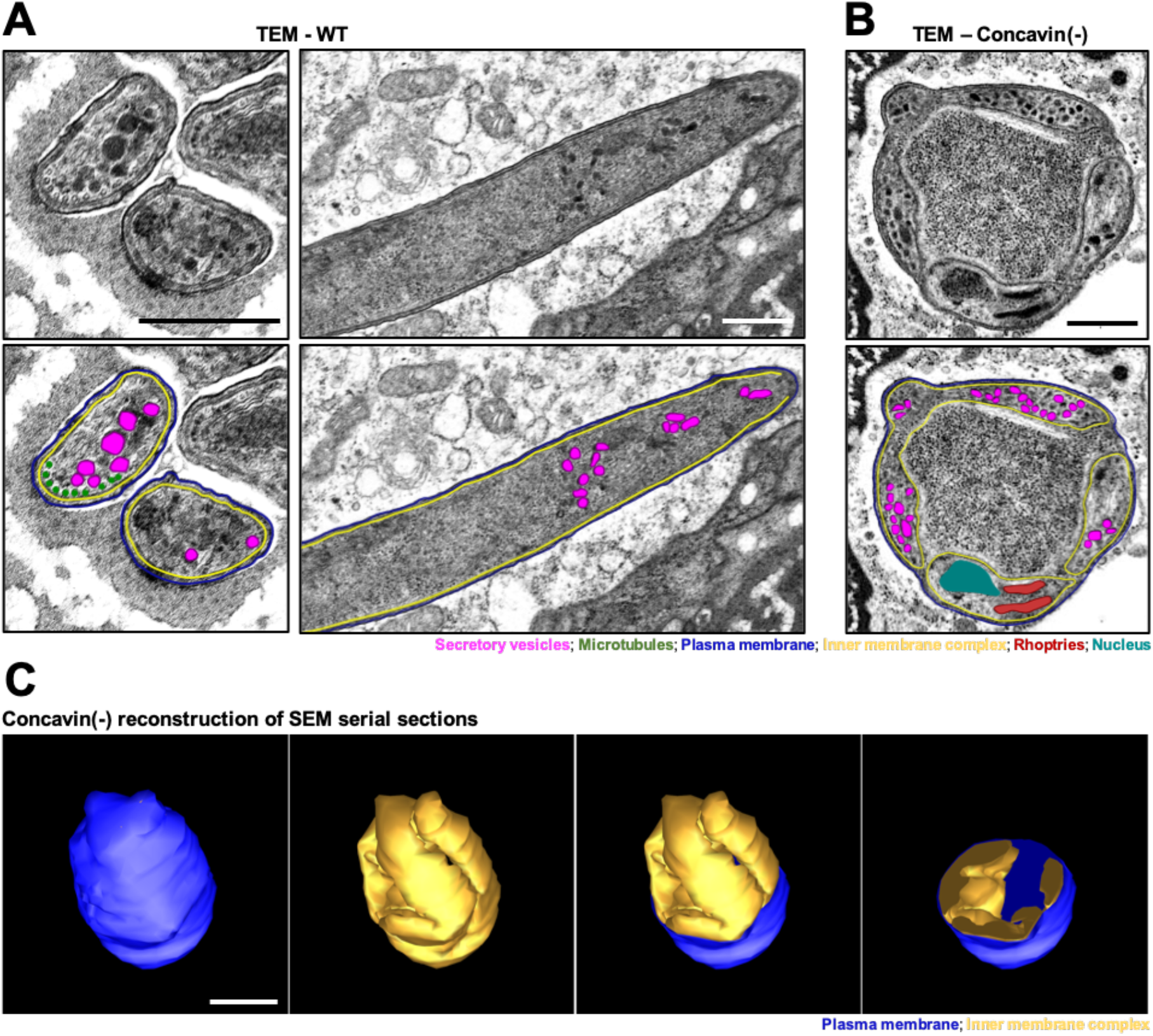
Cytoplasmic IMC extensions in *concavin(−)* sporozoites. **(A)** Transmission electron micrographs of wild type sporozoites with highlighted secretory organelles in magenta, IMC in yellow, microtubules in green and plasma membrane in blue. **(B)** Transmission electron microscopy of *concavin(−)* sporozoites with highlighted secretory organelles as in A; rhoptries in red and nucleus in turquoise. Scale bars 500 nm **(C)** Reconstruction of SEM serial sections of a complete rounded up concavin(−) sporozoite; plasma membrane in blue and IMC in yellow. Scale bar 1μm.

### Complementation with a palmitoylation site mutant partially restores sporozoite form

We next complemented *concavin(−)* parasites with a *concavin* version that expresses an alanine instead of the cysteine of the likely palmitoylation site fused to GFP (Figure 5A, Figure S5). In blood stage parasites concavin^C7A^-GFP displayed a cytoplasmic signal in gametocytes but was absent in asexual stages identical to what we observed in wild type concavin-GFP parasites (Figure 5B). This concavin^C7A^-GFP parasite readily yielded sporozoites in oocysts (Figure 5C). and salivary glands. Quantitation of sporozoite shape from both locations showed that at any time point assessed, over 80% of sporozoites showed a normal shape, even as late as 22 days after infection, when the vast majority of *concavin(−)* sporozoites showed aberrant shapes (Figure 5D, compare to Figure 1D). Only very few sporozoites were completely rounded, while 5-20% of sporozoites showed a rounded proximal end or rounded off around the nucleus. Similar to *concavin(−)*, normal shaped concavin^C7A^-GFP sporozoites were able to move in a wild type manner while deformed sporozoites were only able to move at reduced speed (Figure 5E). This suggests that palmitoylation alone is not essential for concavin function. Furthermore, concavin^C7A^-GFP showed a peripheral localization in both normally shaped and also slightly rounded sporozoites suggesting that localization was also not impaired by the mutation (Figure 5F). The GFP signal in this line appeared similar to the signal obtained by anti-CSP antibodies in *concavin(−)* sporozoites. This suggests that concavin^C7A^-GFP localizes to the plasma membrane and not the IMC.

**Figure 5.**
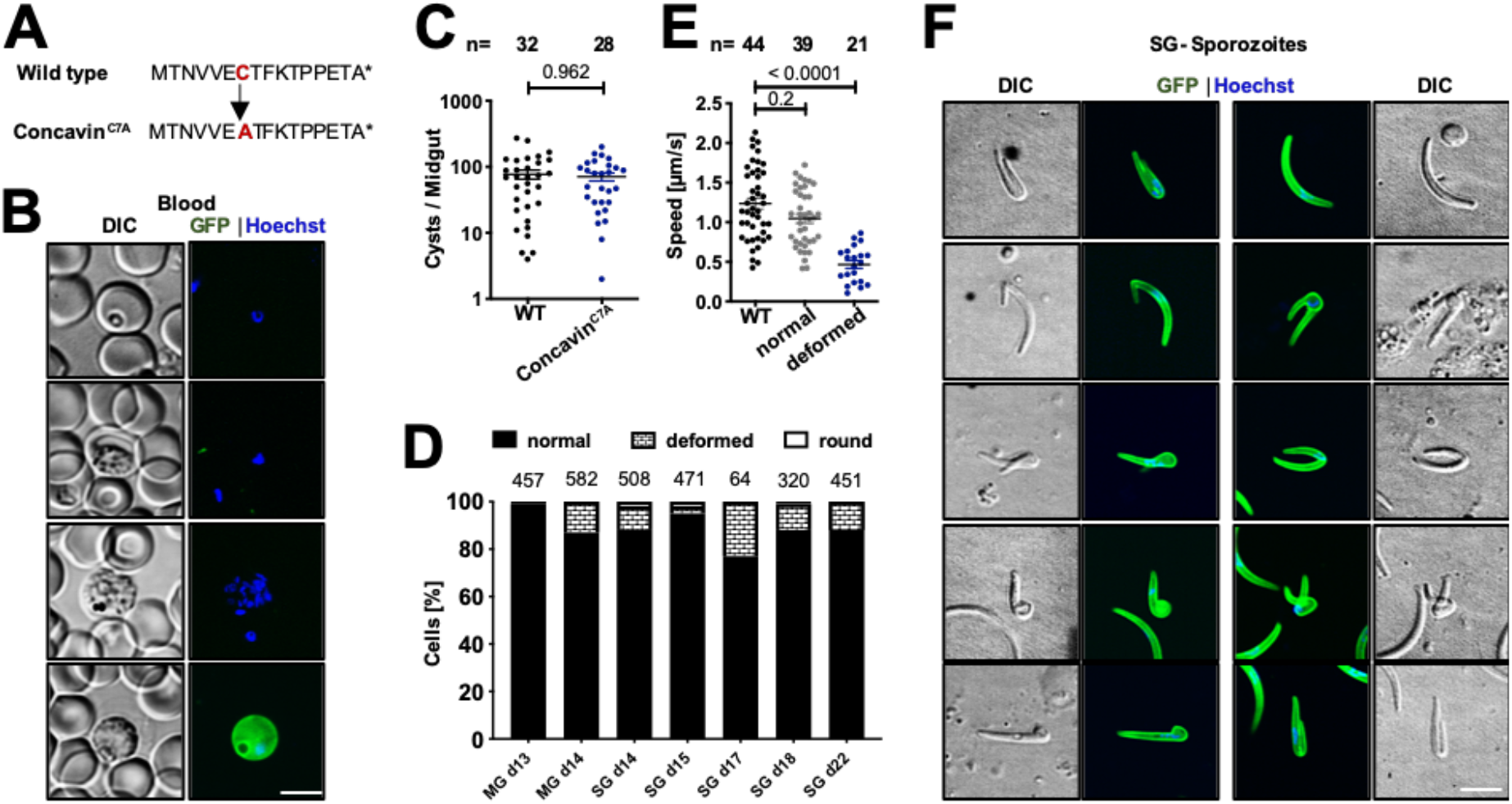
Limited impact of potential palmitoylation for shape maintenance. **(A)** Cysteine 7 is predicted to be palmitoylated and was changed into alanine in the concavin^C7A^-GFP mutant. **(B)** Expression and localization of concavin^C7A^-GFP in blood stage parasites. **(C)** Oocysts number in infected mosquitoes. Data points represent individual midguts observed between d12-17 post infection. Shown is the mean ± SEM. P-value calculated using the Mann Whitney test. **(D)** Quantification of concavin^C7A^-GFP cell shapes at the indicated days from midgut (MG) or salivary gland (SG) derived sporozoites. Numbers above bars indicate investigated sporozoites. **(E)** Average speed of salivary gland sporozoites. Data points represent individual sporozoites. Shown is the mean ± SEM. P-values are calculated using the Kruskal Wallis test followed by the Dunns multiple comparison test. **(F)** Localization of concavin^C7A^-GFP in normal and deformed sporozoites. Scale bar: 5 μm.

### Concavin is essential for efficient transmission

To test if the rounded parasites could still transmit to mice, we let ten infected mosquitoes bite C57BL/6 mice, which are highly sensitive to *P. berghei* infections. All three mice that were bitten by mosquitoes infected with wild type parasites showed the typical blood stage infection in these experiments starting three days after the bites (Figure 6A,B). In contrast, only 8 of 12 mice that were bitten by *concavin(−)* infected mosquitoes ever became infected. In these 8 mice the development of the blood stage infection was delayed by over one day compared to the wild type controls, in itself a loss of infectivity by 90% (Figure 6A,B). To test if deformed sporozoites could enter into cells, we performed an infection experiment, where sporozoites were added to cultured HeLa cells, which are as susceptible to *P. berghei* infection as hepatocytes (Kaiser et al., 2016). After incubation of 1h hour, cells were fixed and labelled with anti-CSP antibodies, without permeabilization, to distinguish sporozoites within and outside of cells. While the first three independent experiments showed a higher percentage of cell invasion for wild type than *concavin(−)* parasites, a fourth experiment lowered the level of statistical confidence (Supplementary Data Table S2). Importantly, however, the deformed parasites could enter into host cells, albeit *concavin(−)* parasites probably entered at an overall lower rate as wild type parasites (Figure 6C). Liver-stage development of *concavin(−)* parasites did not show any difference to wildtype parasites (Figure 6D).

**Figure 6.**
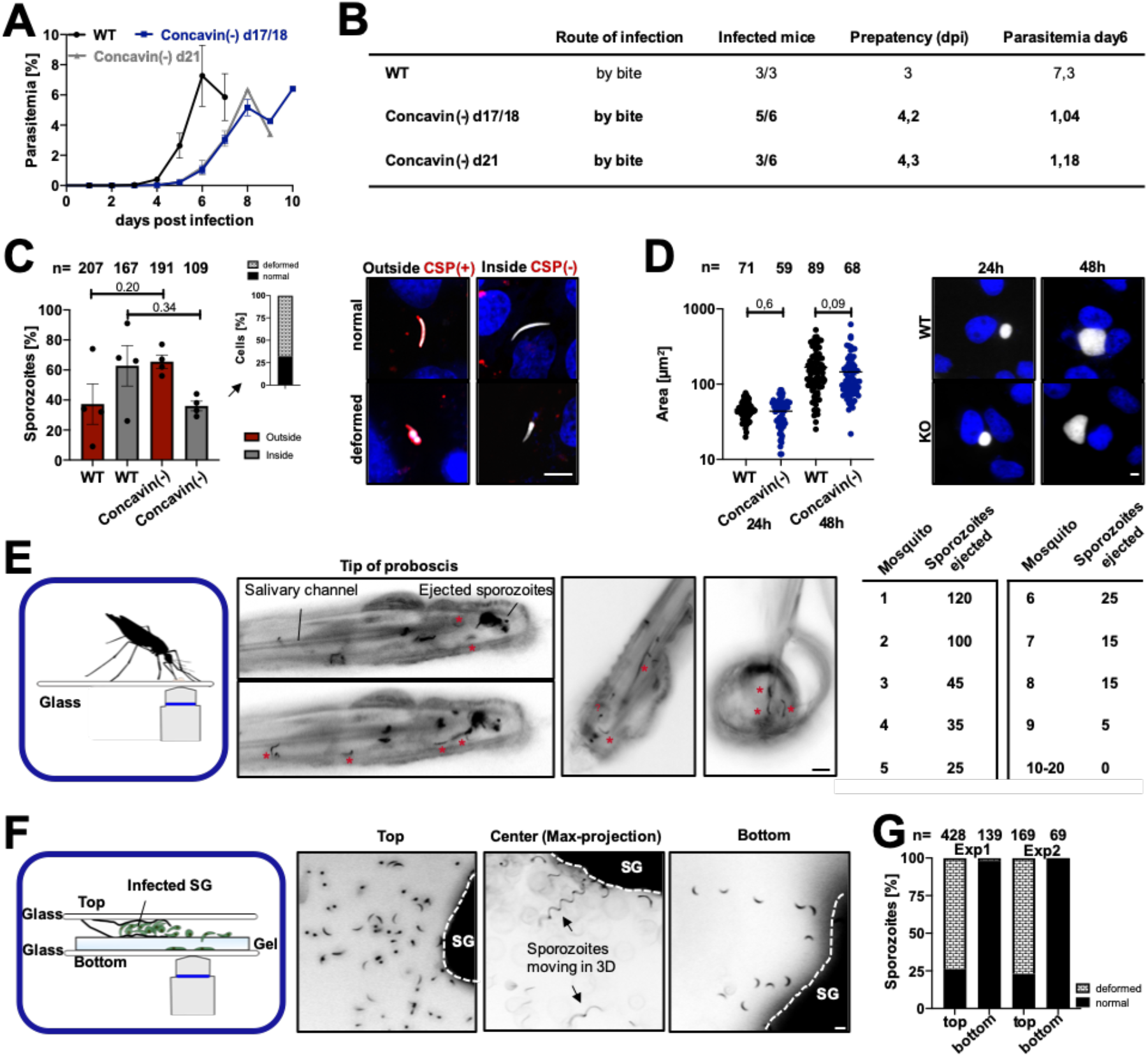
Concavin is essential for efficient transmission by mosquitoes. (**A)** Growth curve of blood stage parasites in C57BL/6 mice infected by the bite of 10 mosquitoes at two different times post infection as indicated. Shown is the mean ± SEM. **(B)** Sum table of infected mice from A with pre-patency period (time to detect a blood stage infection) and parasitemia on day 6. Note that in *concavin(−)* infected mice not all develop a blood stage infection. **(C)** Liver cell invasion assay of *concavin-gfp* and *concavin(−)* parasites. Parasites are positively stained for CSP in case they remain extracellular. Both, normal and deformed parasites were detected intracellularly. Graph shows quantification of CSP positive (red) and negative (grey) sporozoites. Data points represent individual experiments with n indicating the numbers of sporozoites observed. Shown is the mean ± SEM. P-values are calculated using the Mann Whitney test. Scale bar: 5 μm. **(D)** Liver-stage development of *concavin(−)* parasites compared to wildtype, both expressing cytoplasmic GFP (white). Parasite size was measured 24h and 48h post infection. Data points represent individual parasites. Shown is the mean ± SEM. P-values calculated using the Mann Whitney test. Scale bar: 5 μm. **(E)** Sporozoite ejection of immobilized *concavin(−)* infected mosquitoes on glass slides and quantification of ejected sporozoites from 20 mosquitoes. * indicates individual sporozoites in the ejected saliva. Scale bar: 10 μm. **(F, G)** Only normally shaped *concavin(−)* sporozoites released by salivary glands move on helical paths (arrows) through polyacrylamide gels that mimic the skin (F). Quantification of two individual experiments (G): only normal shaped sporozoites were able to migrate through the gel.

But could deformed *concavin(−)* parasites at all reach the liver *in vivo*? Two obstacles might block the progression of the deformed or rounded parasites from the salivary gland to the liver: the narrow salivary ducts through which the parasites are ejected and the dermis in the skin, through which the parasites need to pass prior to entering the blood stream. To test transmission efficiency, we first immobilized *concavin(−)* infected mosquitoes on glass slides and observed the ejection of sporozoites (Figure 7E). As expected (Aleshnick et al., 2020; Frischknecht et al., 2004), ejection of sporozoites was highly irregular (Figure 6E). Yet, at days 17 and 21 in four different experimental sessions we always observed at least one mosquito salivating many dozens of sporozoites while most ejected just a few or none. While not all ejected sporozoites could be readily classified according to their shape the vast majority of sporozoites showed a normal crescent shaped (Movie S3). Some appeared rounded but often revealed themselves as being normally shaped once they reoriented in the focal plane after ejection into the droplet of saliva (Movie S3). We could not confidently see ejected sporozoites that were rounded suggesting that these cannot enter the salivary canal, albeit we cannot exclude that about 10-20% of ejected sporozoites were deformed. Yet, when investigating the sporozoites resident in the salivary gland of the mosquitoes that ejected many sporozoites over 80% of *concavin(−)* sporozoites were deformed. This suggested that rounded sporozoites could not efficiently enter into and pass through the salivary canals. To test if sporozoites fail to migrate through confined spaces, we squeezed a salivary gland between a glass slide and a polyacrylamide gel such that sporozoites were liberated and able to enter into the gel (Figure 6F) (Ripp et al., 2021). On the surface of the gel both deformed and normally shaped parasites are readily visible, while at the bottom end of the gel, only normally shaped sporozoites were found (Figure 6F-G), indicating that deformed sporozoites could not cross the dense matrix of the gel. These data suggest that deformed sporozoites cannot enter and move through confined spaces such as salivary canal and probably also not in skin.

**Figure 7.**
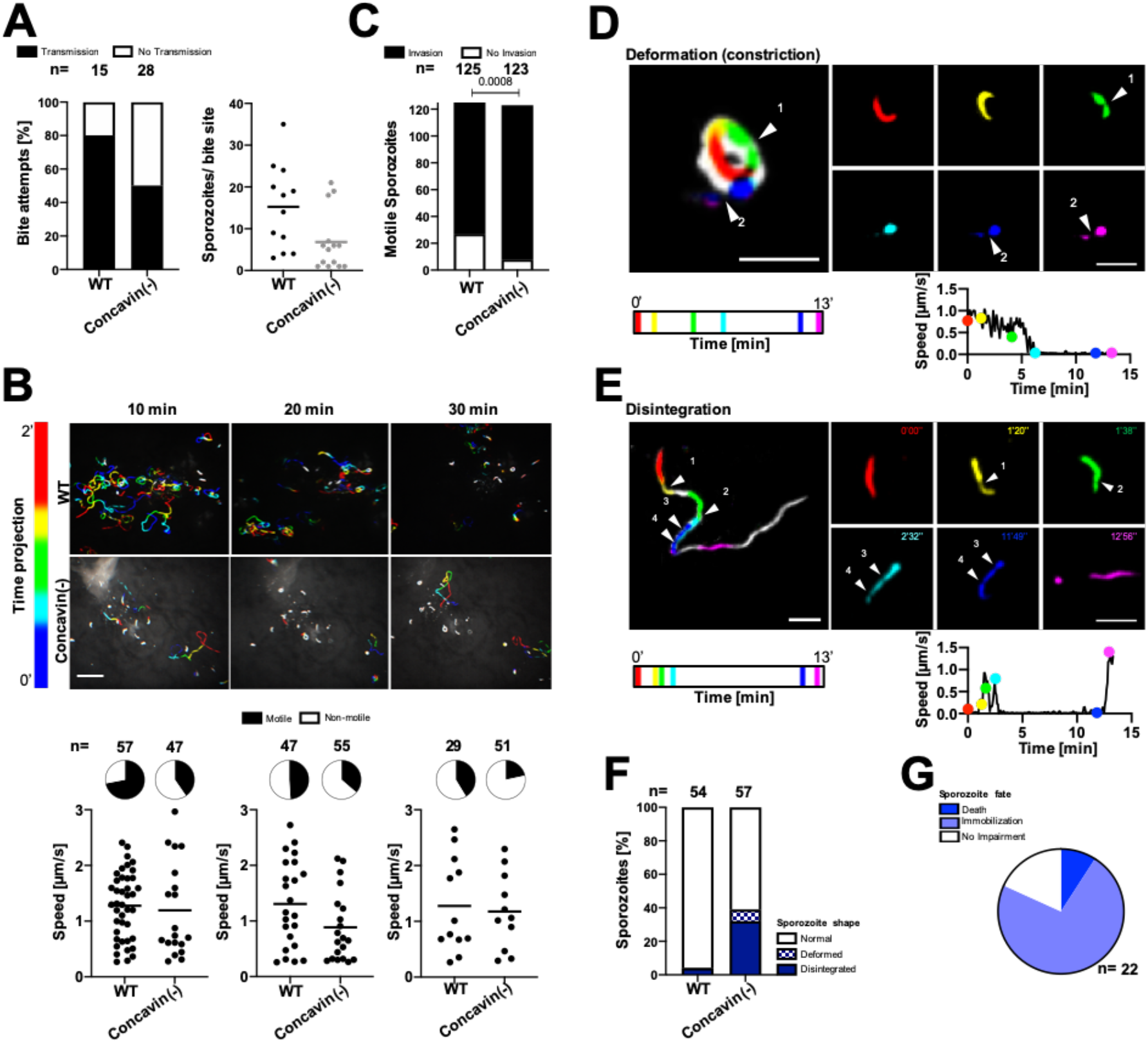
Migrating *concavin(−)* sporozoites lose their cellular integrity. **(A)** Percentage of mosquitoes depositing WT or *concavin(−)* sporozoites in the skin of a mouse during a bite (left) and sporozoites deposited during a mosquito bite (right). **(B)** Maximum fluorescence intensity projections encoded by color for time from movies showing migrating sporozoites after mosquito-bite transmission. Graphs and camembert diagrams below show fraction of motile and immotile sporozoites after 10, 20 and 30 minutes of recording, numbers analysed as well as sporozoite speed. Pooled data from 4-5 WT or 5-8 *concavin (−)* bite sites per time point. Scale bar: 50 μm. **(C)** Motile WT or *concavin(−)* sporozoites entering blood vessels during the first hour following micro-injection. Number above bars shows numbers of sporozoites observed. Pooled data from four WT or five *concavin(−)* experiments. **(D)** Deformation and **(E)** disintegration of *concavin(−)* sporozoites migrating in the skin. Individual images corresponding to the frames shown on the right are indicated in distinct colours in the maximum projection (left) of 13-min movies. Arrowheads indicate constrictions of the parasites. Graphs below time-lapse show that deformation and disintegration are preceded by a decrease in speed. Color of dots correspond to the time-points displayed in the time-lapse images. Scale bars: 10 μm. **(F)** Percentage of deformation and disintegration events observed in 54 WT and 57 *concavin(−)* sporozoites that were tracked for at least 10 min. Pooled data from 5 WT or 10 *concavin(−)* bite sites**. (G)** Consequence of deformation and disintegration from the 22 sporozoites in F include immobilization and parasite death.

### Disintegration of sporozoites migrating in the skin

To investigate if *concavin(−)* sporozoites could progress through the skin, we let mosquitoes infected with wild type or *concavin(−)* sporozoites expressing a cytosolic fluorescent protein bite the ear pinnae of mice and imaged with a spinning disk confocal microscope. This showed that mosquitoes were less successful in transmitting *concavin(−)* than wild type sporozoites. Fewer mosquitoes transmitted *concavin(−)* sporozoites and at lower numbers (Figure 7A), although the lower level of fluorescence in the *concavin(−)* sporozoites might have impeded precise quantification. Intriguingly, of 90 *concavin(−)* sporozoites observed over 13 different bite sites, 72 (80%) appeared to have a normal morphology, suggesting that indeed fewer abnormal sporozoites pass through the mosquito proboscis during a bite (Figure S10A/ B). The remaining 18 (20%) *concavin(−)* sporozoites were abnormally shaped from the beginning of the recording, some being only half as long as normal sporozoites, appearing completely roundish or exhibiting a rounded posterior end (Figure S10A/ B). Analysis of time-lapse series showed fewer migrating mutant sporozoites, which however progressed with the same speed as wild type parasites (Figure 7B; Movie S4). In addition, fewer blood vessel invasion events were observed in motile micro-syringe inoculated *concavin(−)* sporozoites (Figure 7C). Strikingly, some sporozoites formed a bulky dot at their rear (deformation) during migration, mostly associated with passing through a narrow stricture as indicated by their body constriction and decrease in speed (Figure 7D; Movie S5). This round posterior structure sometimes detached as the parasites kept moving forward and this loss of body integrity was defined as disintegration (Figure 7E; Movie S5). Of 54 wild type sporozoites only 2 (4%) showed disintegration, while 32% of 57 migrating *concavin(−)* sporozoites disintegrated and 7% exhibited posterior deformation only (Figure 7F). This suggests that in their effort to migrate in the skin, around 30-40% of *concavin(−)* sporozoites lose their shape or cellular integrity.

Most of those sporozoites stop migrating while some appear to suffer no impairment for the duration of imaging, and around 10% loses viability as evidenced by the loss of GFP fluorescence (Figure 7G). Disintegration most frequently occurred when sporozoites were gliding inside hair follicles, in the upper part of the dermis or after sustained circular gliding, which is usually associated with cell traversal (Formaglio et al., 2014). This suggests that sporozoite disintegration is a consequence of tight interactions between migrating parasites and host cells either because the sporozoites squeeze themselves between or through cells.

Together these data strongly suggest that *concavin(−)* sporozoites fail to maintain their shape, which blocks migration into the salivary ducts and leads to lower levels of transmission. Of the transmitted sporozoites, a large proportion loses their cellular integrity and cannot migrate efficiently in the skin explaining the reduced transmission to the mammalian host.

## DISCUSSION

Here we show that a new protein named concavin contributes to cell shape maintenance in *Plasmodium berghei* sporozoites and is essential for efficient transmission of malaria parasites from mosquitoes to the mammalian host. Although expressed before sporozoites enter salivary glands, the function of concavin only becomes apparent upon prolonged residency within this organ. The parasites appear to rely on concavin to keep the IMC closely subtending the plasma membrane as loss of the protein leads to blebbing of plasma membrane and invaginations of the IMC. The rounding of the sporozoite inhibits ejection from the narrow salivary canals and leads to less efficient deposition of parasites in the skin. Transmitted *concavin(−)* parasites are not yet rounded and can move in the skin as wild type parasites do, but upon squeezing through narrow spaces they lose their cellular integrity much more frequently than wild type parasites. It appears that restraining forces from the environment act upon the motile parasites, which necessitated the evolution of a machinery that keeps the parasites from disintegration. Likely the skin provides a formidable challenge for the parasite as they move for tens of minutes past collagen fibres as well as through and between cells. Based on the number of newly transcribed genes upon salivary gland entry (Matuschewski et al., 2002), the parasite clearly prepares actively for its journey through the skin to the liver. Hence it might be possible that other proteins also contribute to one or multiple complexes that involve concavin and gliding associated proteins for riveting the plasma membrane and IMC.

A strikingly similar loss of cellular integrity during migration of sporozoites is observed in the presence of antibodies targeting CSP (Aliprandini et al., 2018). This surprising similarity shows that the skin is a harmful environment for the moving sporozoite, and indicates that the lack of concavin and the crosslinking of CSP by antibodies can fragilize the parasite. Could these processes be linked? Antibodies crosslink CSP into large aggregates, which are shed at the rear of the parasite and can literally strip a migrating sporozoite. The force acting upon the parasite by these large aggregates being stuck within the tissue as the parasite pushes forward might well lead to a rupture of the plasma membrane-IMC link and hence blebbing. Detailed electron microscopy images of disintegrating parasites would be needed to address if the processes are similar. We postulate that concavin is anchored in the plasma membrane by palmitoylation at its N-terminus, possibly by the palmitoyl-S-acyl transferase DHHC2, which is located at the sporozoite periphery with a bias towards the rear end (Santos et al., 2015). It might hence interact within the membrane with the GPI-anchor of CSP, which could influence the mobility within the membrane of both proteins.

Interestingly, disrupting a predicted palmitoylation site in concavin led to only a very mild phenotypic difference from the wild type with less than 20% of the salivary gland derived concavin^C7A^-GFP parasites showing an aberrant shape. This would suggest that palmitoylation only partially contributes to concavin function and could hint towards the presence of other protein-protein interaction that are essential for concavin functionality and sporozoite shape maintenance. This finding of partial impact of palmitoylation is reminiscent of the recent work on palmitoylation of the myosin light chain in *Toxoplasma gondii* (Rompikuntal et al., 2020), where disruption of palmitoylation led to the partial disassembly of the gliding motor complex but had little impact on motility itself. Considering the EM images showing large invaginations of the IMC away from the plasma membrane in *concavin(−)* sporozoites, we postulate that concavin might contribute towards riveting the plasma membrane to the subtending IMC, which are a constant distance of 25 nm apart. In *T. gondii* the gliding associated protein 45 (GAP45) was shown to bridge the gap between IMC and plasma membrane (Frénal et al., 2010) and fulfil a similar function. GAP45 is associated with the PM through N-terminal palmitoylation and myristoylation and interacts with the IMC at the C- terminus. Intriguingly, *gap45(−) T. gondii* parasites develop normally but show aberrant IMC invaginations upon host cell invasion, reminiscent of the deformed *concavin(−) Plasmodium* sporozoites (Egarter et al., 2014; Frénal et al., 2010). Strikingly, concavin is the first protein essential for sporozoite shape maintenance that is not fixed to or appears as part of a cytoskeletal structure. Compared to *Plasmodium berghei* GAP45 which consists of 184 aa, concavin is about twice as large (393 aa) and hence might well span the distance between plasma membrane and IMC. How the protein contributes to keeping the IMC in place remains elusive. Understanding how it remains mobile while contributing to cellular shape maintenance clearly needs further work. The presence of a potential palmitoylation site at the C- terminus of concavin, in combination with the ability of the protein to recover after FRAP suggest an incorporation into the PM rather than the IMC. The ability of normal and deformed sporozoites to remain motile excludes most likely a function of concavin in glideosome formation. We speculate that concavin is either directly involved in PM and IMC organisation or through transient interactions with GAP45. Open questions include: does concavin interact with other proteins and if yes, what are the binding partners? Does it interact with proteins linked to the IMC or the actin-myosin motor machinery that drives the parasite? Likely more proteins are important in keeping the plasma membrane at a constant distance from the IMC and maintaining the shape of these parasites.

Sporozoites are not born with a final shape, but mature from long and slender ones in the oocysts to crescent-shaped slightly thicker ones in the salivary glands (Kudryashev et al., 2012; Muthinja et al., 2017). Investigations by cryogenic electron tomography revealed that the sub-pellicular network (SPN) is only robustly detectable in sporozoites isolated from the salivary glands (Kudryashev et al., 2012) and tagging of SPN proteins revealed their peripheral localization only in salivary gland derived sporozoites (Khater et al., 2004). This suggests that the SPN and possibly its linkage to microtubules plays a key role in generation of the crescent shape observed for transmission ready sporozoites. In turn, this suggests that the shape has a key function for sporozoite infectivity. Indeed, mutants where the shape of sporozoites is altered have been shown to transmit less efficiently or not at all to rodent hosts (Montagna et al., 2012; Spreng et al., 2019; Tremp et al., 2013; Volkmann et al., 2012). Yet, *concavin(−)* sporozoites reveal for the first time a loss of cellular integrity as a phenotypic consequence of deleting a *Plasmodium* gene. The movies of sporozoites migrating in the mouse skin *in vivo* suggest that plasma membrane containing considerable amounts of cytosol is lost as the parasites migrate through tight strictures.

Upon liver cell invasion, the sporozoite naturally changes its shape, a likely active process depending on newly translated proteins from stored transcripts (Gomes-Santos et al., 2011). Deletion of the RNA binding protein pumilio-2 led to a progressive rounding of sporozoite already in the salivary glands and premature expression of liver stage specific genes (Gomes-Santos et al., 2011; Lindner, Mikolajczak, et al., 2013). In contrast to pumilio-2 mutants, which only round up after several days of salivary gland residence, the lack of concavin led to early rounding after salivary gland invasion (Figure 1C-E). This, and the different localization of the two proteins suggests a completely different function of the two proteins. Also, *pumilio-2(−)* sporozoites ‘round up’ in a way reminiscent of the shape changes after liver cell invasion: the sporozoites bleb in the center with their ends initially keeping their sporozoite shape (Gomes-Santos et al., 2011). This is similar to the rounding of mutants lacking the IMC-1 and IMC-1h proteins (Khater et al., 2004; Volkmann et al., 2012). In contrast *concavin(−)* sporozoites round up from their proximal ends. (Figure 1B, 5C-E and Figure S8).

We found that in *T. gondii* disrupting the concavin orthologue has no visible impact on *in vitro* life in cultured fibroblasts. This however does not rule out that during other parts of the life cycle of this parasite more constraining barriers encountered by the parasite might impact on cellular integrity too. We observed a slight reduction in oocyst numbers, which hints at a possible function of concaving also for ookinetes. Ookinetes need to pass the peritrophic matrix that forms around the ingested blood meal and through one layer of epithelial cells. Ookinetes move much slower than sporozoites and hence concavin might not be as important during their short life as it is for the much longer living and faster migrating sporozoites facing many more constrictions on their journey from oocyst to liver.

In conclusion, we showed here the disintegration of migrating *Plasmodium* sporozoites due to the lack of a novel protein, concavin. Concavin was identified by a proteomics analyses of secretion. Functional analyses through GFP-tagging, FRAP, gene deletion and *in vivo* imaging showed the importance of concavin for the maintenance and integrity of *Plasmodium* sporozoite shape and hence efficient transmission from mosquito to mammal.

## MATERIALS AND METHODS

### Generation of parasite lines

#### Concavin(−)

The 3’UTR (779 bp) of PbANKA_1422900 was amplified from wild type gDNA using primers JK57 and JK58 and inserted into a plasmid (pL22) containing the recyclable yFCU/ hDHFR selection cassette and *gfp* expressed under the *hsp70* promoter digested with NotI and SacII. The 5’UTR (554 bp) was amplified using primers JK55 and JK56 and inserted into the plasmid using KpnI and HindIII. The resulting plasmid pL24 was linearized with KpnI and SacII prior transfection for double crossover integration (Figure S3; Table S1).

#### Marker free concavin(−) NS

The drinking water of mice infected with *concavin(−)* parasites was supplemented with 2 mg/ml 5-FC (5-fluorocytosine). Clonal parasites which looped out the selection cassette were obtained by limiting dilution. (Figure S3; Table S1)

#### P. berghei complementation

The 5’UTR together with the entire ORF of PbANKA_1422900 was amplified from wild type gDNA using primers JK55 and JK176 and inserted into a plasmid (pL59) containing *gfp* and the *TgDHFR* selection cassette using KpnI and NdeI. Resulting in plasmid pL79. For transfection of *concavin(−)NS* parasites via double crossover the plasmid was digested with KpnI and SacII. (Figure S5; Table S2)

#### PF3D7 complementation

The *P. berghei* 5’UTR was amplified using primers JK55 and JK179 and inserted into pL59 using KpnI and BstbI. Followed by the insertion of PF3D7_0814600 amplified with primers JK177 and JK178 from P. *falciparum* gDNA and digested with BstbI and NdeI. For transfection of *concavin(−)NS* parasites via double crossover the plasmid was digested with KpnI and SacII. (Figure S5; Table S2)

#### Concavin^C7A^ complementation

*Concavin^C7A^* together with *gfp* was amplified from pl79 using primers JK236 and JK237. The resulting PCR product was digested with BamHI and ligated into pL82 digested with BamHI and BstBI with filled in overhangs. Leading to the final plasmid pL120. For transfection of *concavin(−)NS* parasites via double crossover the plasmid was digested with KpnI and SacII. (Figure S5; Table S2)

#### Phil1-GFP

PhIL1-GFP (PBANKA_020460) parasites with GFP at the C-terminus were generated via single homologous recombination. A region of the PhIL1 gene 54 bp downstream of the ATG start codon and lacking the stop codon was amplified using primers P969 and P970 (Figure S6; Table S2). The PCR product was digested using EcoR1 and BamH1 enzymes and ligated into a vector containing the *TgDHFR* selection cassette as a positive selection marker. For transfection the final vector was linearized using BsaB1.

Transfection of linearized plasmids was performed as previously described (Janse et al., 2006). After electroporation of schizonts, positive selection was performed using pyrimethamine. All resulting monoclonal lines were obtained through limiting dilution using NMRI or CD1 mice. Briefly, 0,9 parasites were injected into 10 naïve mice followed by genotyping of positive animals on day 9 post infection.

### Mosquito infections

All experiments were performed with *Anopheles stephensi* mosquitoes, fed with 1% salt/water and 10% sucrose/water solution containing 0.05% Para- aminobenzoic acid (PABA). Naïve mosquitoes were kept at 28°C and 70% humidity and subsequently transferred to 21°C after infection. Infection of mosquitoes was done with mice infected with 20 million blood stage parasites, 4 days post infection. Gametocyte formation was monitored by counting exflagellation events in peripheral blood. Mice were anesthetized with a combination of ketamin/ xylazin and mosquitoes were allowed to bite for 20 min.

### Sporozoite motility assay

Gliding assays of isolated sporozoites were performed in 96-well optical bottom plates (Nunc) using 3% BSA/ RPMI with a frame rate of 3 seconds for 3 minutes on a Zeiss CellObserver widefield microscope with 25 x magnification. Speed was determined with the manual tracking tool in ImageJ.

### Immunofluorescence staining of sporozoites

Sporozoites were seeded with 3% BSA/ RPMI into an 8well labtek chambered cover glass. After fixation with 4% PFA for 20 minutes, cells were permeabilized with 0.5% TritonX for 15 minutes. Primary antibodies were incubated for 1 hour and washed twice with PBS. Secondary antibodies together with Hoechst were incubated for 1 hour. Cells were washed twice with PBS and observed under the microscope. Images were either taken on a Zeiss CellObserver widefield (63x) or Nikon/PerkinElmer spinning disc (100x) microscope. Image processing was performed with ImageJ. Antibodies: rabbit anti GFP for IFA 1/40 (abfinity 0,4 μg/ μl), mouse anti CSP (mAb 3D11, 1/100, Yoshida et al 1980), goat anti-mouse Alexa 594 1/1000 (Invitrogen 2 mg/ml), goat anti-rabbit Alexa 594 1/1000 (Invitrogen 2 mg/ml). For STED imaging anti-mouse Atto 594 1/300 (Sigma) and anti-rabbit Atto 647N 1/300 (Sigma) was used. Staining of sporozoites with antibodies or Sir tubulin was performed as described previously (Spreng et al., 2019)

### FRAP

FRAP experiments were performed using a PerkinElmer Nikon spinning disc microscope with FRAP module. Images were taken with a 100x objective (NA 1,4) every 0,5 seconds. Selected regions were bleached using a 405 nm laser at 100% laser power.

### STED imaging

Super- resolution imaging was performed on a STED microscope from Abberior Instruments GmbH (Göttingen) using a 100x objective. STED- illumination was done using a 594 nm or 640 nm excitation laser in combination with 775 nm depletion. Images were taken with 15 nm pixel size and line accumulation of 2. Deconvolution was done using the Imspector software provided by Abberior using the Richardson- Lucy algorithm with 30 iterations. Further image processing was done using Fiji.

### Electron Microscopy

Isolated salivary glands from infected *Anopheles stephensi* mosquitoes were directly dissected into 2% glutaraldehyde and 2% paraformaldehyde in 100 mM Cacodylate buffer and fixed at 4°C overnight. After rinsing in buffer the samples were further fixed in 1% osmium in 100 mM Cacodylate buffer for 1 hour, washed in water, and incubated in 1% uranylacetate in water overnight at 4°C. Dehydration was done in 10 minute steps in an acetone gradient followed by stepwise Spurr resin infiltration at room temperature and polymerization at 60°C.

The blocks were trimmed to get cross sections of the salivary glands and sectioned using a Leica UC6 ultramicrotome (Leica Microsystems Vienna) in 70 nm thin sections. The sections were placed on formvar coated slot grids, post-stained in uranyl acetate and Reynold‘s lead citrate and imaged on a JEOL JEM-1400 electron microscope (JEOL, Tokyo) operating at 80 kV and equipped with a 4K TemCam F416 camera (Tietz Video and Image Processing Systems GmBH, Gautig).

### Array tomography

3D reconstruction of the round sporozoites was done using array tomography. Ultrathin serial sections (70 nm) were cut on an UC7 ultramicrotome (Leica Microsystems, Vienna, Austria) equipped with a section receiver (cd-fh, Heidelberg) enabling a smooth pick up of serial sections on wafers (Si-Mat SiliconMaterials) treated by glow discharge. The sections were post-stained by placing the wafer pieces in uranyl acetate 20 min and lead citrate for 10 min, each in a closed tube. The wafers with sections were washed repeatedly in water after each step. Serial sections through one sporozoite (around 40 sections) were imaged by scanning electron microscopy with a LEO Gemini 1530 equipped with a field emission gun and an ATLAS scanning generator (Zeiss). Images of 10.000 x 10.000 pixels and 2nm resolution were taken at the same area in consecutive sections using the in-lens detector. Imaging parameters used: 3 mm working distance, 30 μm aperture and 2 kV acceleration voltage. The images through each parasite were aligned and the membranes segmented manually using the IMOD software package (Kremer et al, 1996). In total 6 sporozoites were reconstructed.

### Sporozoite invasion into cells and liver- stage development

Hela cells were seeded into 8 well Labtek chamber slides with glass bottom. Once the cells reached about 95% confluency sporozoites were added on top of the cells. The entire slide was centrifuged for 5 minutes at 800 rpm to make the sporozoites adhere to the cells and incubated at 37°C. After 1 hour the cells were fixed with 4% PFA/ PBS for 30 minutes. Followed by staining of the cells against CSP (mouse anti CSP mAb 3D11, 1/ 100, Yoshida et al 1980). The primary antibody was incubated for 1 hour and washed twice with PBS. The secondary antibody (goat anti-mouse Alexa 594 1/1000 (Invitrogen 2mg/ml)) together with Hoechst were incubated for 1 hour. Cells were washed twice with PBS and observed under the microscope. Images were either taken on a Zeiss CellObserver widefield (63x) or Nikon/PerkinElmer spinning disc (100x) microscope. Image processing was performed with Fiji.

For liver-stage development cells were split into 2 wells 1 hour post infection and fixed with 4% PFA/ PBS after 24 h and 48 h. Nuclei were stained with Hoechst and images taken on a Zeiss Axio-Observer widefield (63x) microscope. Image processing and quantification of parasite size was done with Fiji.

### Live imaging of Mosquito sporozoite ejection

For imaging of ejected sporozoites mosquitoes were immobilized with small drops of superglue on the wings and thorax on a 24x 60 mm cover glass. The labrum was carefully removed using 2 needles to liberate the stylets of the mosquito which were ideally pressed flat onto the cover glass. Imaging was done on a Zeiss Axio- Observer with 25x Objective at a frame rate of 3 frames per second.

### 3D Imaging in polyacrylamide gels

Polyacrylamide hydrogels were prepared as described before (Pelham & Wang, 1997). Briefly, APS and TEMED were added to a prepolymer solution containing 3% acrylamide and 0.03% bis-acrylamide in PBS. The solution was pipetted onto a silanized glass coverslip immediately and covered with a second glass coverslip. After polymerization, the top coverslip was removed in PBS. Whole salivary glands were dissected into 30 μl 3% BSA/RPMI and covered with the hydrogel. Images were recorded on top of the hydrogel as well as at the hydrogel-glass interface using a 25x objective and a frame rate of 3s.

### *In vivo* imaging of sporozoites in the mouse skin

*In vivo* imaging in the skin was performed on a spinning-disk confocal system (UltraView ERS, Perkin Elmer) controlled by Volocity (Perkin Elmer) and composed of 4 Diode Pumped Solid State Lasers (excitation wavelengths: 405 nm, 488 nm, 561 nm and 640 nm), a Yokogawa Confocal Scanner Unit CSU22, a Z-axis piezoelectric actuator and a Hamamatsu Orca-Flash 4.0 camera mounted on a Axiovert 200 microscope (Zeiss). Z-stacks of 6 plans covering 25 to 30 μm were acquired using a LCI “Plan- Neofluar” 25x/0.8 Imm Korr DIC objective (Zeiss) at a rate of 2.7 frames per second for up to 80 minutes following sporozoite transmission.

For bite transmission experiments, infected *Anopheles stephensi* were selected under an epifluorescence stereomicroscope 14-15 days after the infectious blood meal and used between day 18 to 21 post-infection. To enhance bite rate, mosquitoes were deprived of sucrose one day before the experiment. For intravital imaging of syringe-inoculated sporozoites, infected salivary glands were harvested in 1x DPBS 13 to 21 days following the infectious blood meal and kept intact on ice. Shortly before the experiment, they were crushed, filtered on a 35-μm strainer and further diluted in 1x DPBS. All experiments were performed using either concavin KO sporozoites or a control *P. berghei* ANKA strain expressing GFP under the control of the *hsp70* promoter (Ishino et al., 2006).

Prior to imaging, 4- to 6-week old female C57BL/6JRj mice (Janvier Labs) were injected intravenously into the tail vein with 10-15 μg of Alexa Fluor™ 647-coupled anti-CD31 antibody (clone 390, Biolegend) to label blood vessels. Animals were then anesthetized with a mixture of ketamine (125 mg/kg body weight, Imalgene 1000, Merial) and xylazine (12,5 mg/kg body weight, Rompun 2%, Bayer), and their ear pinnae were gently epilated with a piece of tape. The dorsal side of their ears was then either exposed to mosquito bites for two minutes or injected with 0.2 μL of sporozoite suspension using a micro syringe (NanoFil 10 μL syringe mounted with a 35G beveled needle, World Precision Instruments), yielding fields of view containing in average 100 parasites. Animals were then immediately transferred onto the microscope stage to localize and image the inoculation sites. During acquisition, mice were kept warm with a heating blanket (Harvard Apparatus) and their anesthesia status was regularly monitored (Amino et al., 2007).

Image files were processed and quantified using Fiji (Schindelin et al., 2012). Parasite morphology in the skin was determined on the first images obtained immediately after localization of the bite site. Sporozoite movements were manually tracked over 2-min movies recorded at the indicated time points after inoculation and mean velocity was determined using the MTrackJ plug-in (Meijering et al., 2012). Sporozoites whose speed was inferior to 0.25 μm/s were considered immotile. Blood vessel invasion events were quantified over the first hour following sporozoite micro-injection and normalized to the number of motile sporozoites in the field at 10 min post-infection. To quantify the frequency of parasite disintegration after bite transmission, only sporozoites which could be tracked for at least 10 min were taken into consideration. Following loss of parasite integrity, gradual disappearance of fluorescence was considered a hallmark of sporozoite death.

### *T. gondii* parasite culture

Parasites were cultured in Human Foreskin Fibroblasts (HFFs) using Dulbecco’s modified Eagle’s medium (DMEM) supplemented with 10% fetal calf serum, 2mM L-glutamine and 10mg/mL gentamycin. Cell culture was maintained at 37°C and 5% CO2.

### *T. gondii* parasite strain generation

#### Parasites with TGGT1_216650 endogenously tagged and floxed

Endogenous tagging was done as described in (Stortz et al., 2019). Briefly, RHΔku80DiCre tachyzoites (created by Dr Moritz Treeck (Hunt et al., 2019) were used for endogenous tagging. Guide RNAs to target both the C-terminal region, and upstream of the predicted 5’UTR of TGGT1_216650 were designed. These guides were cloned into a plasmid which expressed Cas9-YFP. The 5’ loxP was introduced into the parasites as an oligo flanked by 33bp of sequences homologous to the region that the Cas9 targeted. The repair templates used for the introduction of the C-terminal tags (YFP and SNAP) were amplified by PCR (Q5 polymerase, New England Biolabs) from template plasmids. The primers used for the amplification of these tags were designed in such a way that an LIC sequence was used as a linker between the protein and the tags, whereas a loxP sequence was introduced downstream of the tags. Homology sequences 50bp long were added on either side of the tags to facilitate homologous recombination. The 5’loxP and 3’ tags were introduced into the parasites in two separate successive transfections. 10μg of vector (encoding the guide RNA and Cas9-YFP) and the respective repair templates were transfected into 1×107 newly egressed parasites, using 4D AMAXA electroporation. Following transfection, the parasites were allowed to invade and replicate for approximately 40hrs, at which point they were mechanically egressed, filtered, and sorted for transient Cas9-YFP expression using a cell sorter (FACSAria IIIu, BD Biosciences) into 96-well plates. Successfully genetically modified parasite clones were identified by IFA, PCR and sequencing (using primers designed to bind as indicated by the blue and red arrows in Figure S4.

### Induction of TGGT1_216650 knock-out

RHΔku80DiCre tachyzoites with a floxed TGGT1_216650 C-terminally tagged with a SNAP-tag were cultured in the presence of 50nM rapamycin for 1 week and then cloned out by serial dilution in a 96-well plate. Successful knockouts were identified by PCR and sequencing using primers designed to bind as indicated by the black arrows in Figure S4.

### *T. gondii* Plaque assay

5×10^2^ parasites were used to inoculate 6-well plates in the presence of dimethyl sulfoxide (DMSO) as vehicle control or 50nM rapamycin. After 7 days of undisturbed incubation, the cells were fixed with 100% ice cold methanol for 20mins at room temperature, and washed with phosphate buffered saline. The cells were then left in eosin for 1min, followed by 2mins in methylene blue, and finally washed thoroughly with water.

### *T. gondii* Immunofluorescence assays and microscopy

HFFs were inoculated with parasites for 24hrs, after which they were either imaged live or fixed with 4% paraformaldehyde (PFA) for 20mins at room temperature. For live imaging, the dyes (SNAP-Cell 647-SiR, and HALO-tag Oregon Green) were used as per the manufacturer’s instructions. For fixed samples, the cells were washed 3 times with phosphate buffered saline (PBS) following fixation, and then permeabilized and blocked for 45mins at room temperature with 3% bovine serum albumin (BSA), 0.2% Triton X-100 in PBS. The cells were then labelled with the following primary antibodies for 1hr at room temperature; mouse anti-GFP (1:500, monoclonal Anti-GFP, Roche), rabbit anti-GFP (1:1000, polyclonal Anti-GFP, abcam), rabbit anti-GAP45 (1:1000, a generous gift from Dominique Soldati Favre; (Plattner et al., 2008)), mouse anti-IMC1 (1:2000, a generous gift from Gary Ward; (Tilley et al., 2014)). After incubating with primary antibodies, the cells were washed 3 times with PBS and then labelled with the following secondary antibodies for 1hr at room temperature in the dark; STAR 635P anti-rabbit (Abberior), STAR 635P anti-mouse (Abberior), Alexa Fluor488 anti-mouse (Life Technologies), Alexa Fluor Plus 488 anti-rabbit (Invitrogen). Following the incubation with the secondary antibodies, the cells were incubated with 0.4μM Hoechst for 5mins, and finally washed 3 times with PBS and mounted with ProLongTM Gold antifade mounting medium (ThermoFisher Scientific). Widefield images were taken using a 100x objective on a Leica DMi8 widefield microscope attached to a Leica DFC9000 GTC camera. Z-stacks were taken and then processed with ImageJ.

### Ethics statement

All animal experiments were performed according to European guidelines and regulations and the German Animal Welfare Act (Tierschutzgesetz) and executed following the guidelines of the Society of Laboratory Animal Science (GV-SOLAS) and of the Federation of European Laboratory Animal Science Associations (FELASA). All experiments were approved by the responsible German authorities (Regierungspräsidium Karlsruhe) or Animal Care and Use ommittee of Institut Pasteur (CETEA 2013-0093) and the French Ministry of Higher Education and Research (MESR 01324).

### Animal work

For all experiments female 4-6-week-old Naval Medical Research Institute (NMRI) mice, Swiss or C57BL/6 mice obtained from Charles River or Janvier laboratories were used. Transgenic parasites were generated in the *Plasmodium berghei* ANKA background (Vincke & Bafort, 1968) either directly in wild type or from wild type derived. Parasites were cultivated in NMRI or Swiss CD1 mice while transmission experiments with sporozoites were performed in C57Bl/6 mice only. Animal experiments conducted at the Institut Pasteur were approved by the Animal Care and Use Committee of Institut Pasteur (CETEA 2013-0093) and the French Ministry of Higher Education and Research (MESR 01324) and were performed in accordance with European guidelines and regulations.

### Statistical analysis

Statistical analysis was performed using GraphPad Prism 8.0 (GraphPad, San Diego, CA, USA). Data sets were either tested with a Mann Whitney or Kruskal Wallis test. A value of p<0.05 was considered significant.

## Acknowledgements

We thank Miriam Reinig and rotating student helpers and the team of the Centre for the Production and Infection of Anopheles (CEPIA, Institut Pasteur) for rearing *Anopheles stephensi* mosquitoes as well as Markus Ganter for helpful discussions and comments on the manuscript. We acknowledge the microscopy support from the Infectious Diseases Imaging Platform (IDIP) at the Center for Integrative Infectious Disease Research and the expert help from the Mass Spectrometry and Proteomics Core Facility at the Center for Molecular Biology (ZMBH) of Heidelberg University.

## Author contributions

J.K. and F.F. designed the project; J.K., P.F. C.F. J.M.M., D.B., C.L., J.R., J.G. and F.F. performed research; all authors analyzed data; M.M. and R.A. supervised the *Toxoplasma* and *in vivo* imaging parts, respectively. J.K. and F.F. wrote the paper with input from all authors.

## SUPPLEMENTARY INFORMATION

**Supplementary Figure S1.**
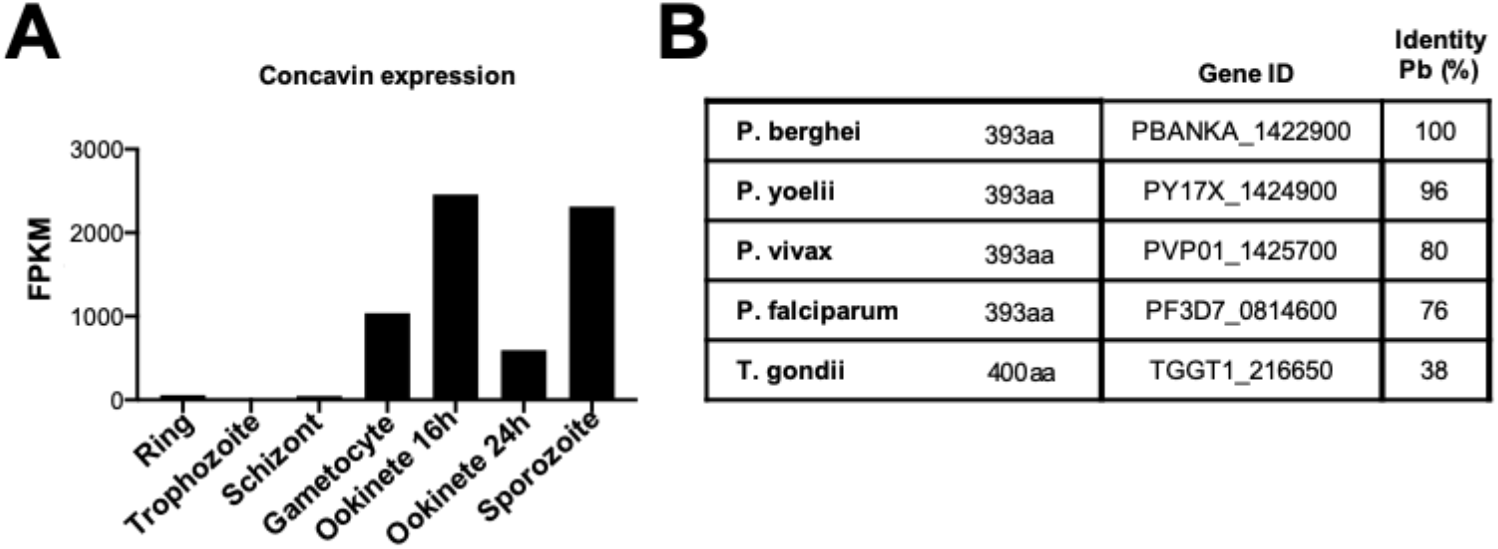
**(A)** RNAseq abundance of concavin in blood and mosquito stage parasites **(B)** Sequence identity of *P. berghei* concavin with *P. yoelii*, *P. vivax*, *P. falciparum* and *T. gondii*.

**Supplementary Figure S2.**
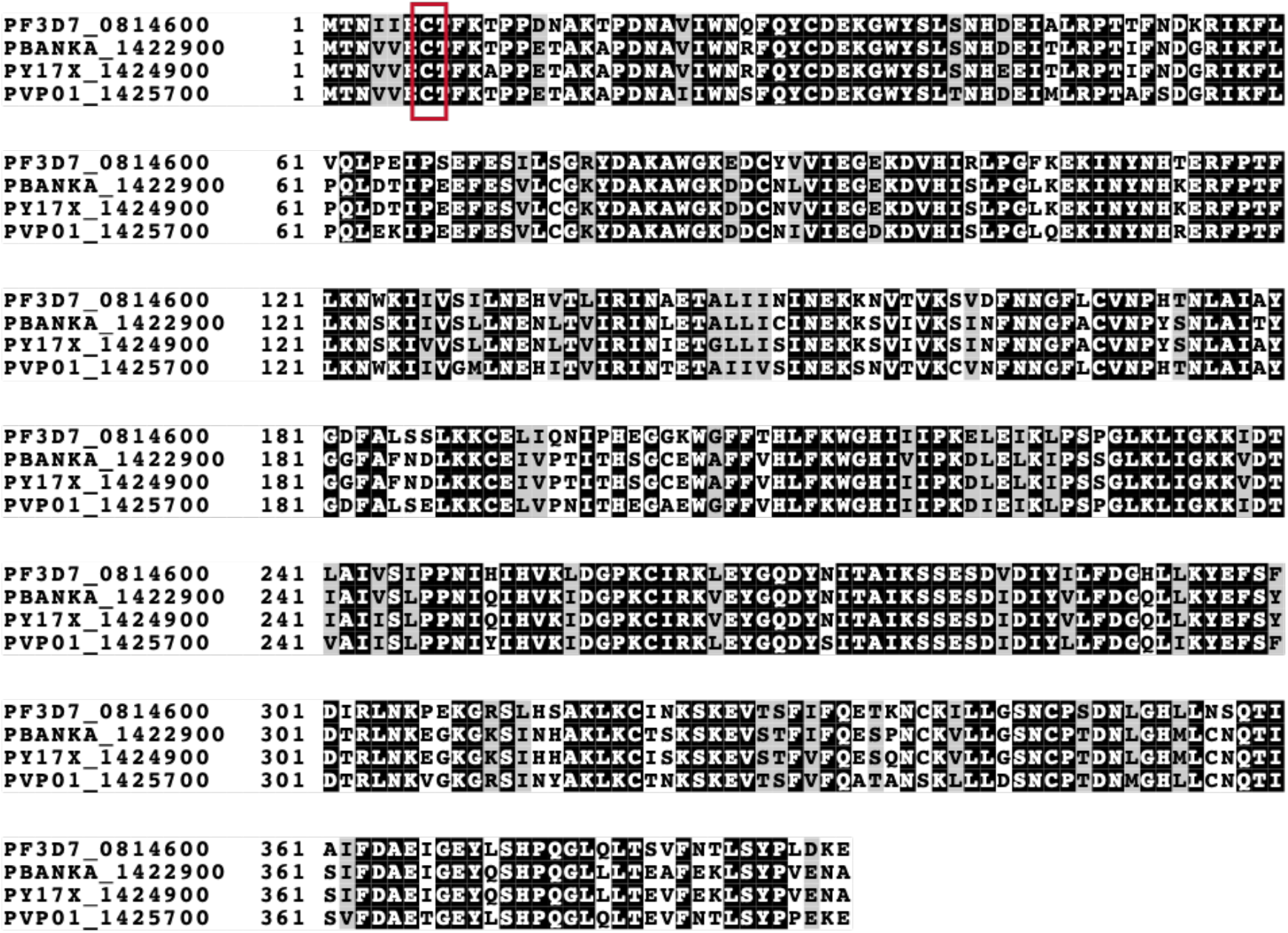
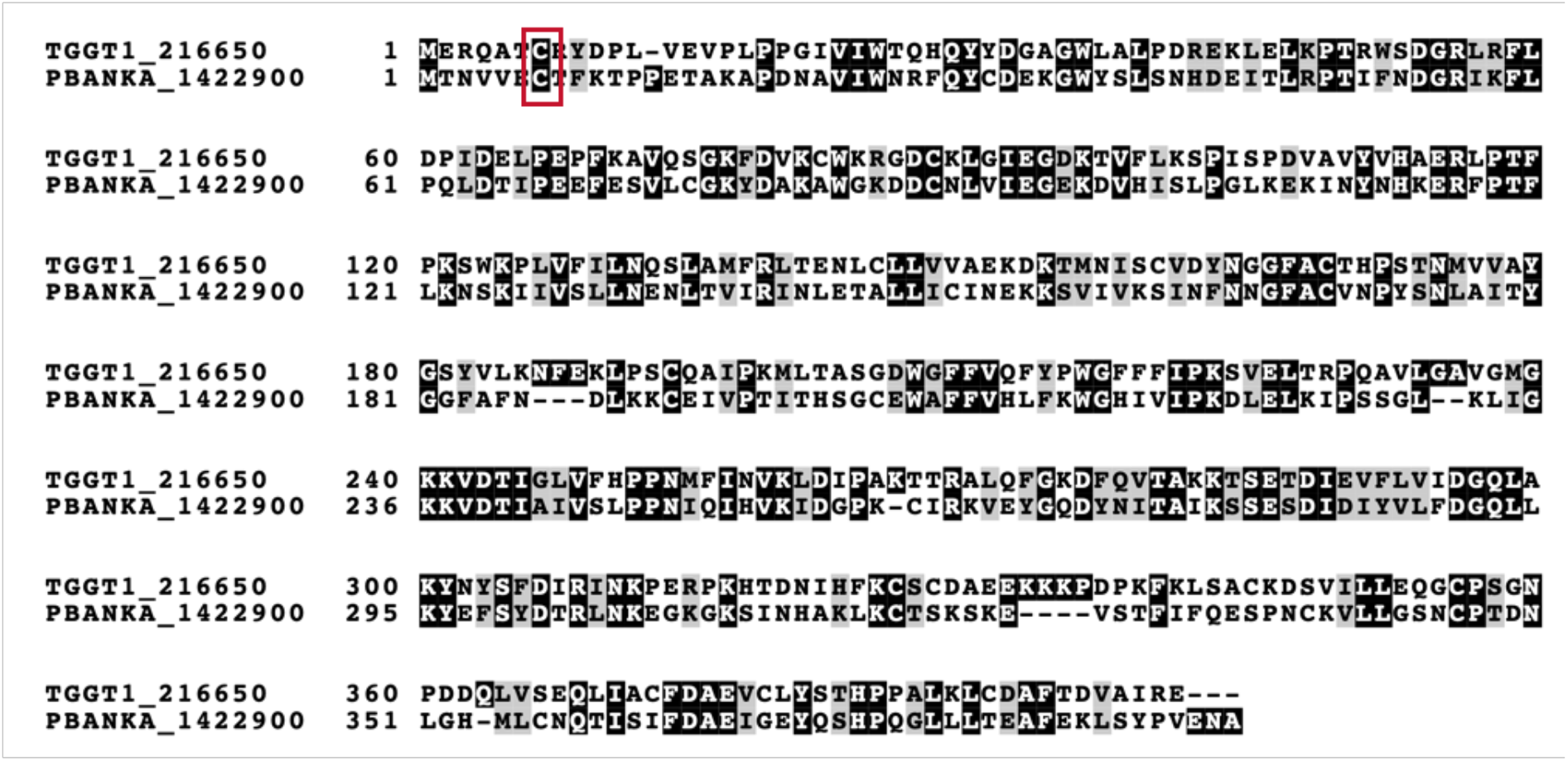
**(A)** Clustal Omega Multiple sequence alignment of *Plasmodium spp*. **(B)** Clustal Omega Multiple sequence alignment with the *T. gondii* orthologe. Potential C- terminal palmitoylation site is highlighted in red.

**Supplementary Figure S3.**
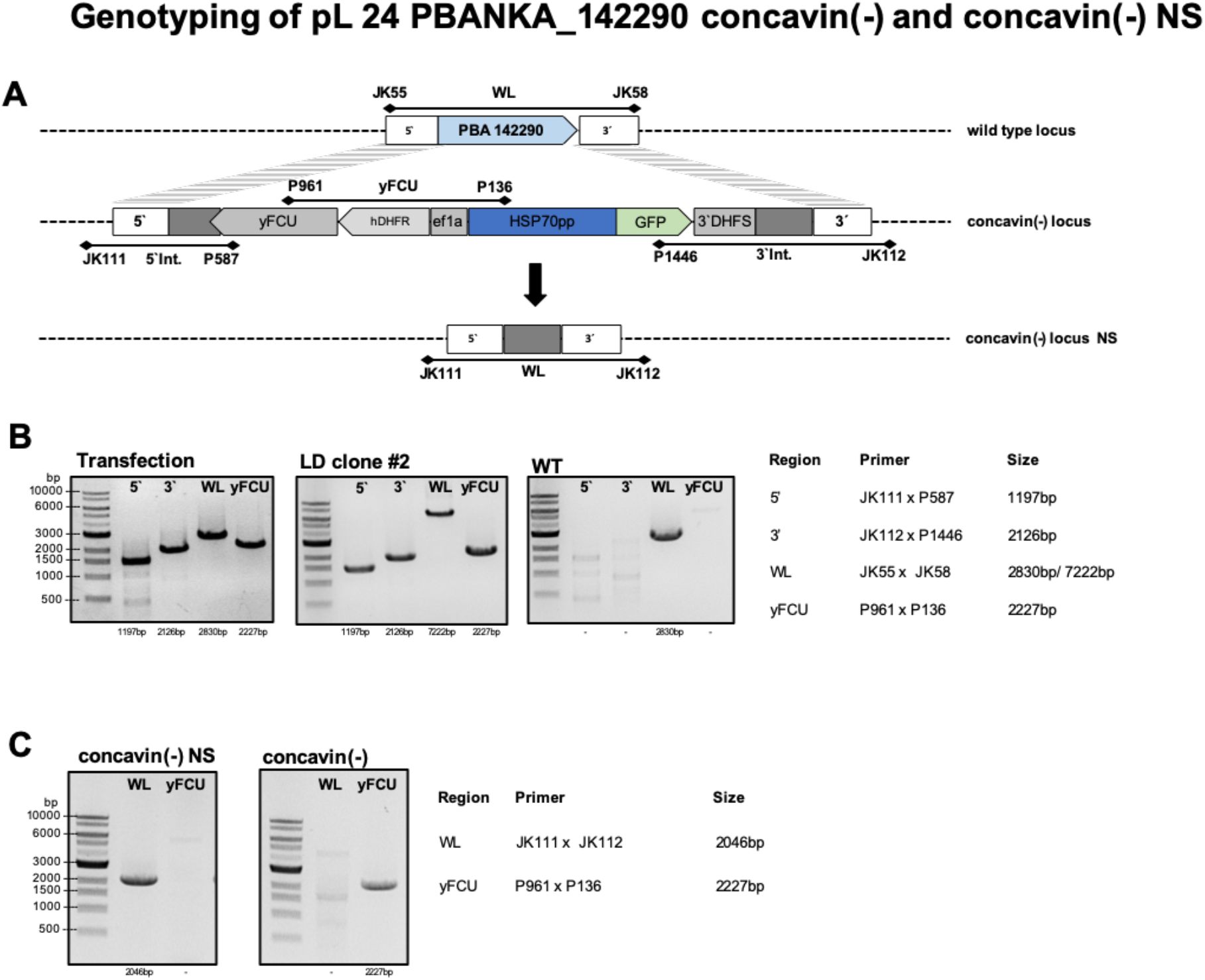
Generation of *concavin(−)* and *concavin(−) NS* parasites via double homologous recombination. **(A)** Cartoon showing the cloning strategy and primers used for genotyping. **(B)** Genotyping PCRs of non-clonal *concavin(−)* parasites directly after transfection and after limiting dilution. Agarose gel pictures show 5’integration, 3’integration as well as wildtype and selection marker as indicated in A. Expected amplicon sizes are indicated on the right. **(C)** Genotyping PCRs of *concavin(−)* parasites after looping out the selection cassette. Expected amplicon sizes are indicated on the right.

**Supplementary Figure S4.**
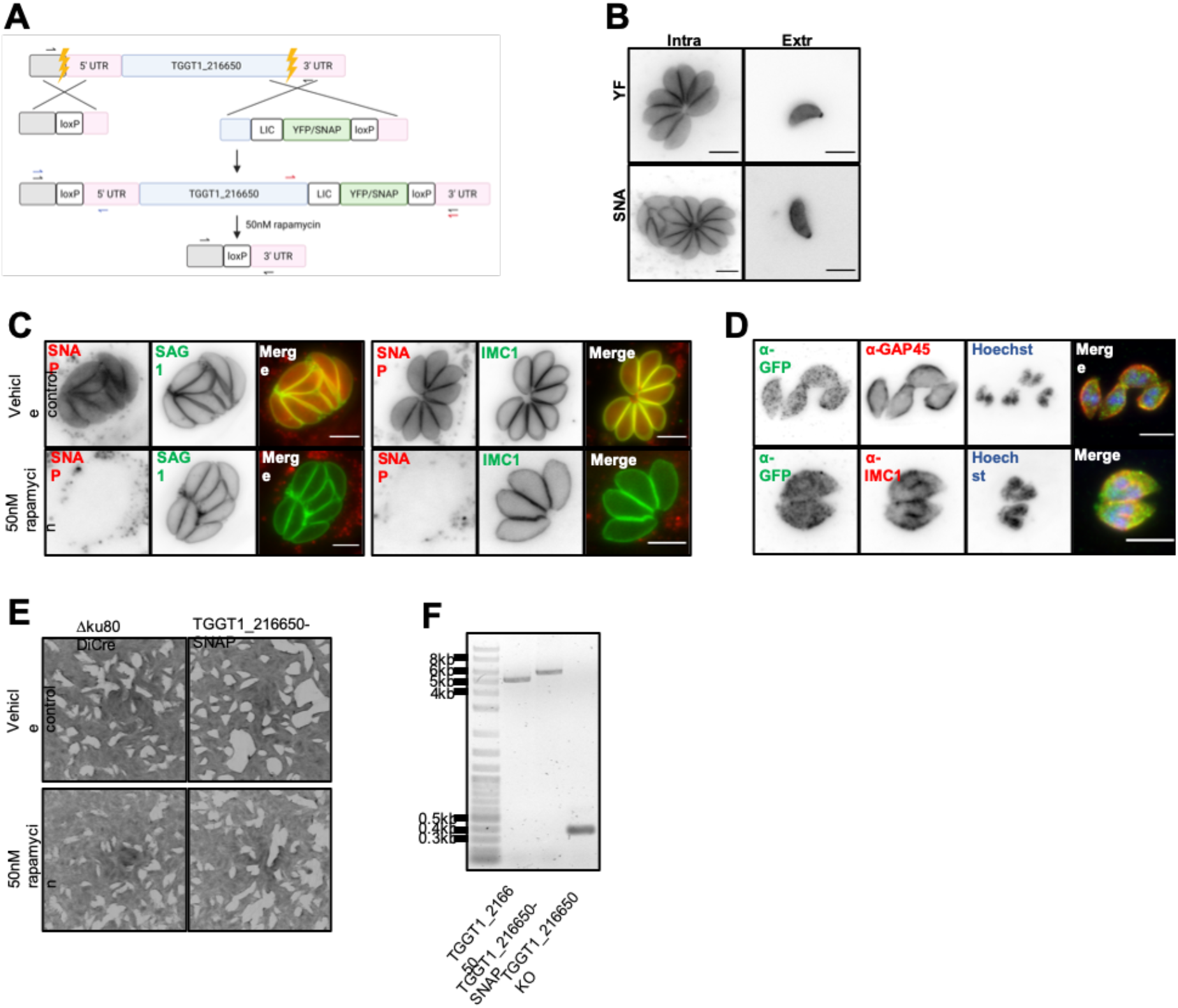
TGGT1_216650 is non-essential in *T. gondii* tachyzoites. **(A)** CRISPR/Cas9 was used to induce double strand breaks. The DNA repair templates used were designed with homology arms to favour homologous recombination. The approximate position of the 5’ UTR was estimated based on the TGME49_216650 annotation on ToxoDB. The LIC sequence was used as a linker between the gene and the tags. Correct integration was confirmed via PCR and sequencing, the primer binding sites as indicated with red and blue arrows. Upon addition of 50 nM rapamycin, the Cre recombinase subunits expressed in the parasite strain dimerise, excising the floxed sequence. **(B)** Images show the localisation of TGGT1-216650 endogenously tagged with YFP and SNAP tags. Parasites were imaged live in both intracellular and extracellular conditions. **(C)** SAG1 and IMC1 were internally and C-terminally tagged with HALO-tag respectively using the same protocol as in panel A. Upon knockout of TGGT1_216650, no phenotype was observed. The parasites were imaged live. **(D)** The parasites were fixed and antibodies were used to amplify the signal. The gene of interest was not observed at the daughter cells while still inside the mother cell during division. **(E)** 7-day plaque assays. Knockout of the gene of interest following addition of 50 nM rapamycin had no effect on the fitness of the parasites. **(F)** A knockout line was successfully obtained and can be maintained in culture. Confirmation of successful knockout via both PCR and sequencing, the primer binding sites as indicated with black arrows in panel A. All scale bars: 5 μm.

**Supplementary Figure S5.**
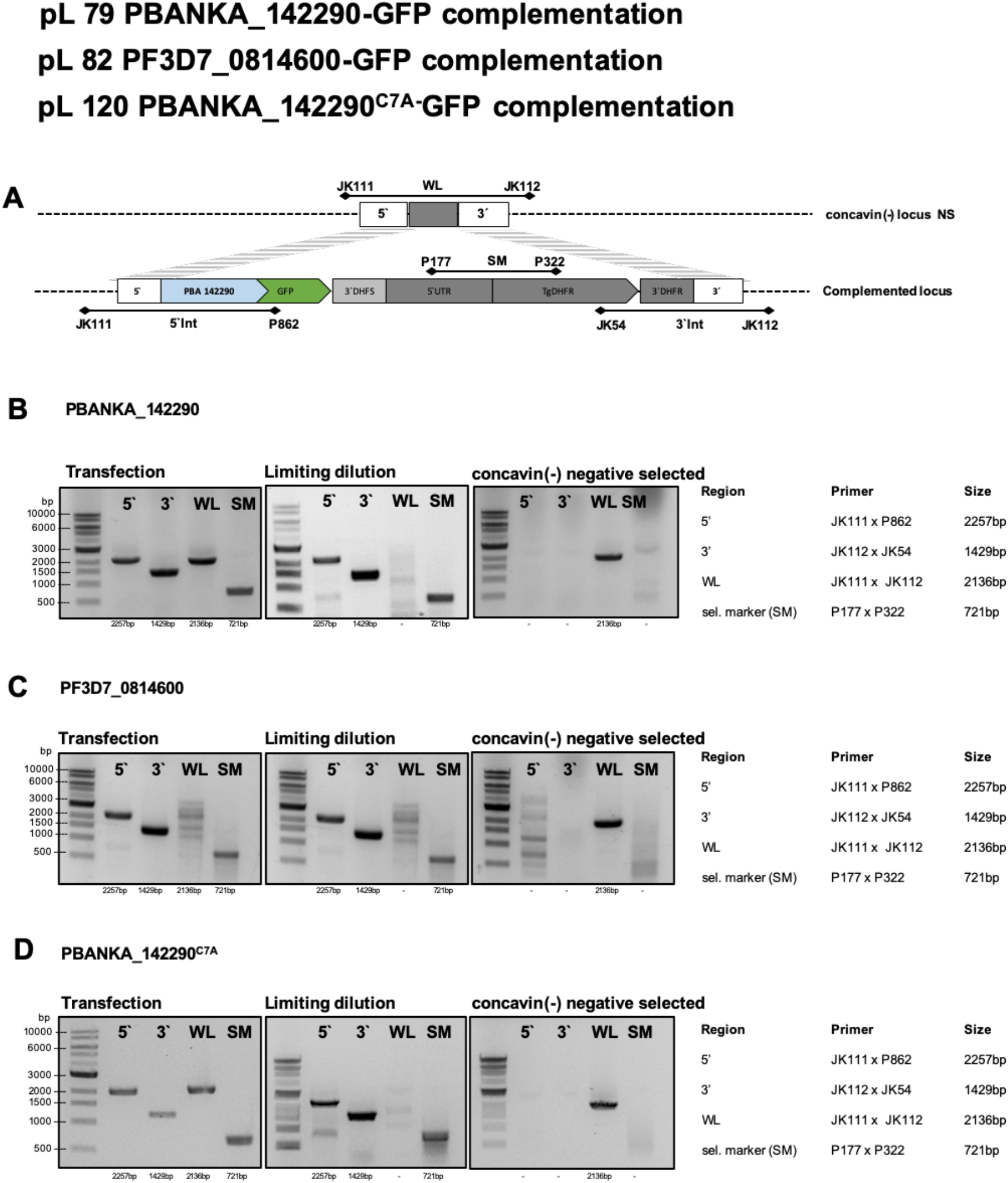
Generation of *concavin(−)|P.berghei-gfp*, *concavin(−)|P.falciparum-gfp* and *concavin*^C7A^ parasites via double homologous recombination into *concavin(−)* NS parasites. **(A)** Cartoon showing the cloning strategy and primers used for genotyping. **(B)** Genotyping PCRs of the non-clonal *concavin(−)|P.berghei-gfp* parasite line directly after transfection and after limiting dilution. Agarose gel pictures show 5’integration, 3’integration as well as wildtype and selection marker as indicated in A. Expected amplicon sizes are indicated on the right. **(C)** Genotyping PCRs of the non-clonal *concavin(−)|P.falciparum-gfp* parasite line directly after transfection and after limiting dilution. Agarose gel pictures show 5’integration, 3’integration as well as wildtype and selection marker as indicated in A. Expected amplicon sizes are indicated on the right. **(D)** Genotyping PCRs of the non-clonal *concavin^C7A^* parasite line directly after transfection and after limiting dilution. Agarose gel pictures show 5’integration, 3’integration as well as wildtype and selection marker as indicated in A. Expected amplicon sizes are indicated on the right.

**Supplementary Figure S6.**
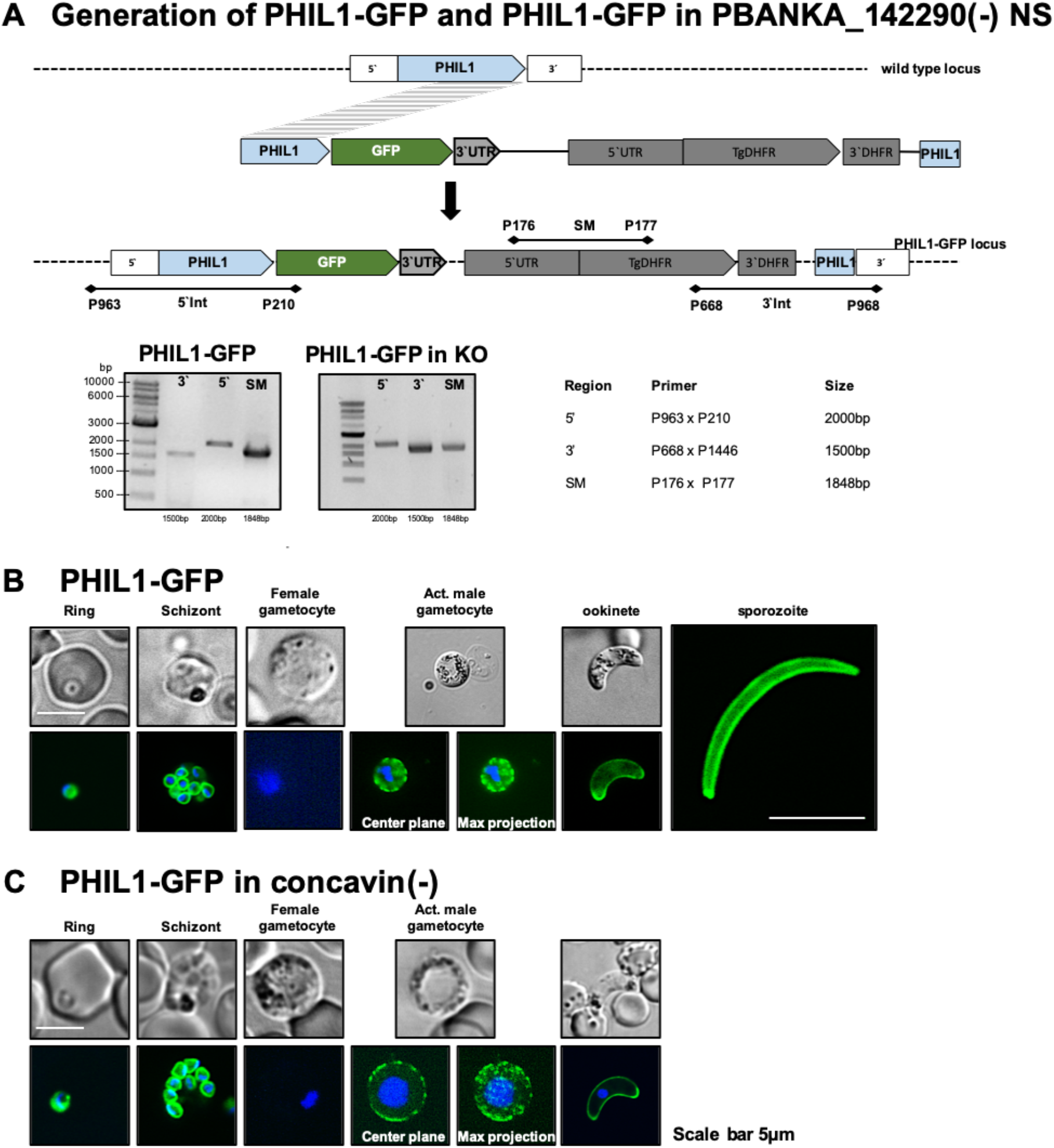
**(A)** Generation of *phil1-gfp* and *concavin(−)|phil1-gfp* parasites via single homologous recombination. The cartoon shows the cloning strategy and primers used for genotyping. Agarose gel pictures shows genotyping of the non-clonal parasites in the wild type as well as in the *concavin(−)* background. (B) Localization of PhiL1-GFP (green) in wild type parasites. Nuclei (blue) are stained with Hoechst. Scale bar: 5 μm. **(C)** Localization of PhiL1-GFP (green) in *concavin(−)* parasites. Nuclei (blue) are stained with Hoechst. Scale bar: 5 μm.

**Supplementary Figure S7.**
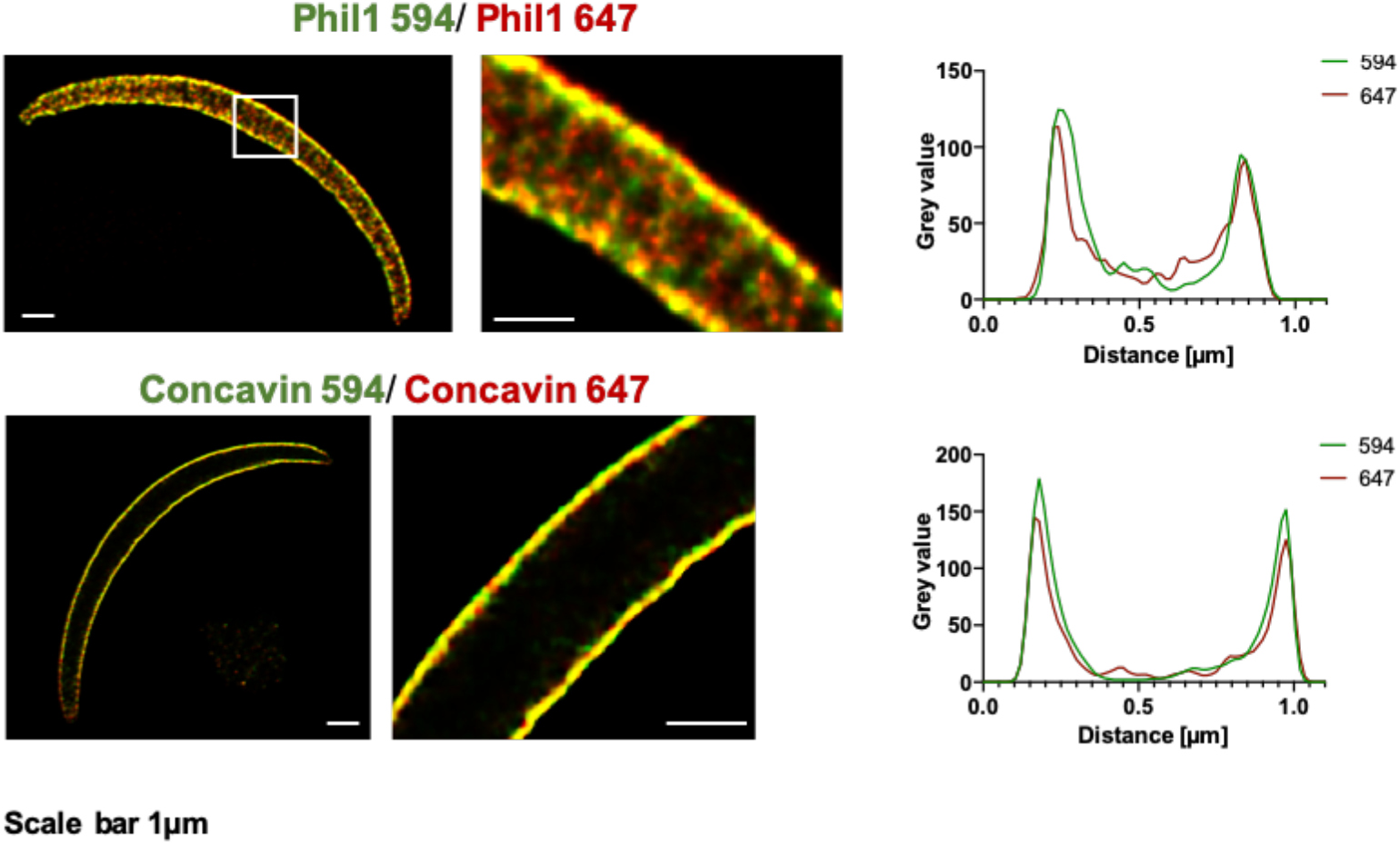
Control images used for STED. Bleed through of signal in cells stained with atto-594 into the 647 channel resulted in overlays with almost no difference in distance between the 2 channels. Images were deconvolved using the Richardson-Lucy algorithm. The distance between the 2 signals was measured using the plot profile of the respective channels in Fiji (Figure 3C). Measurements and plot profiles taken at the center of the cell.

**Supplementary Figure S8.**
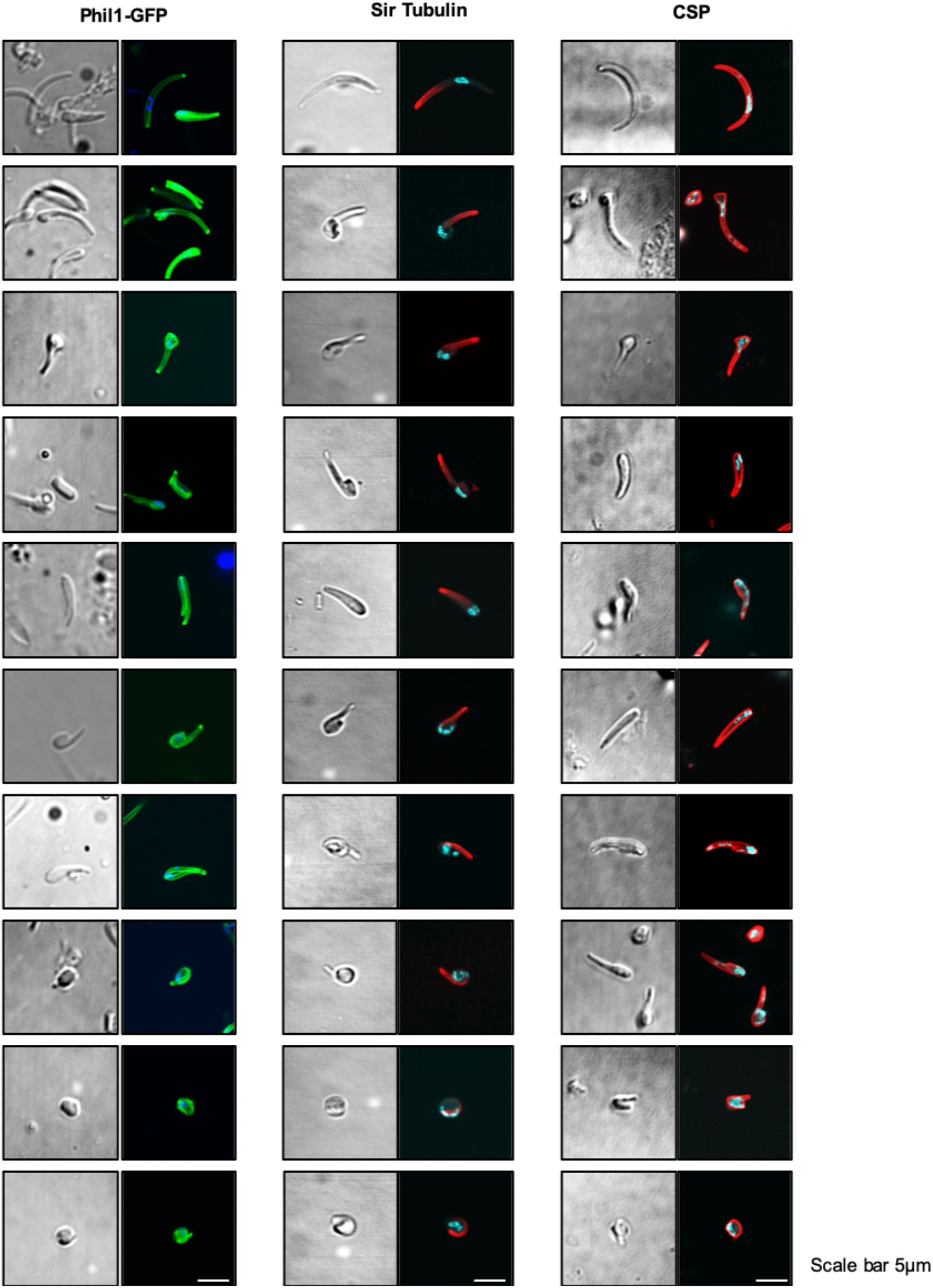
Image gallery of *concavin(−)* parasites expressing *phil1-gfp* (green) or stained with SiR-tubulin or anti-CSP antibody (red). Scale bar:

**Supplementary Figure S9.**
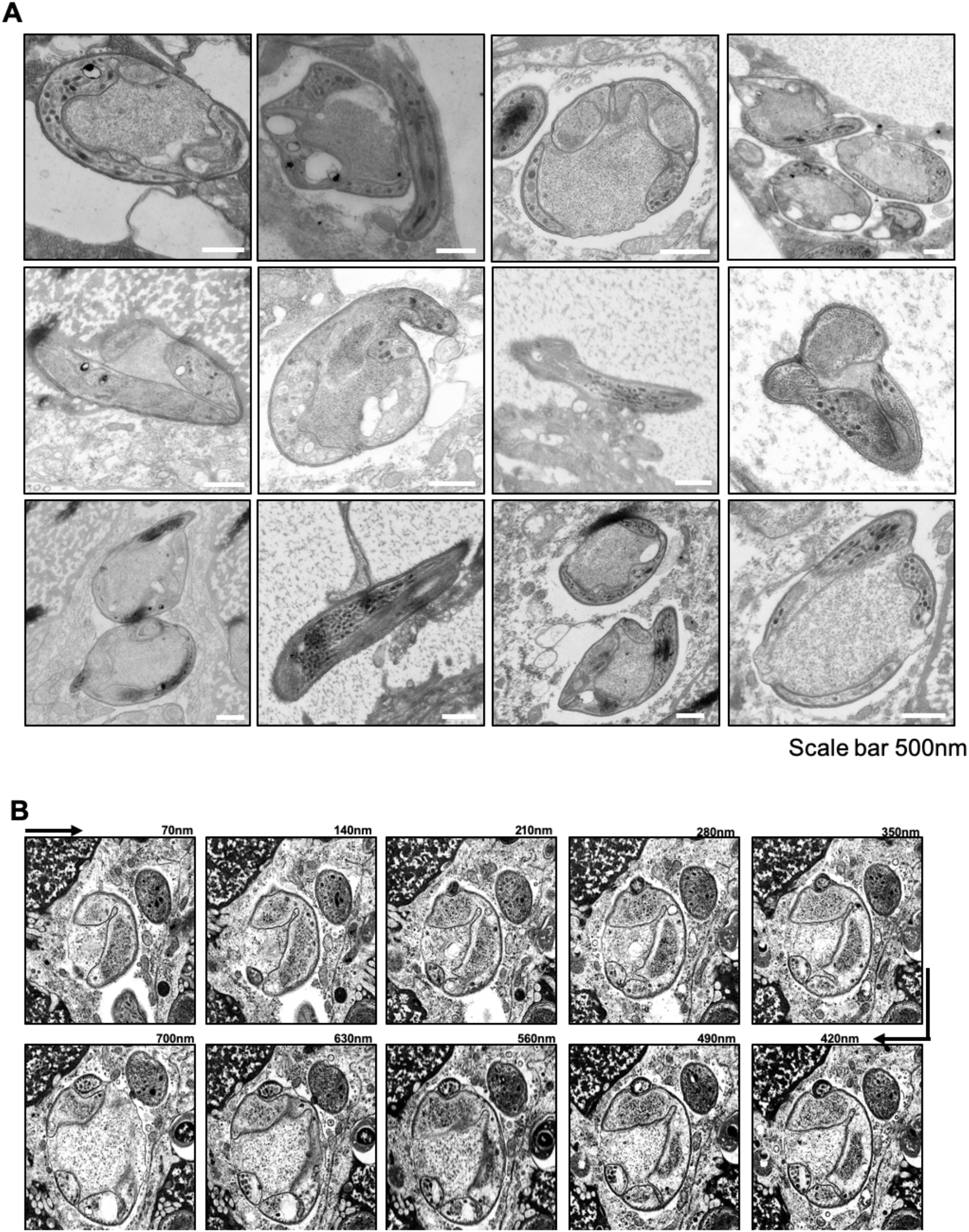
**(A)** TEM image gallery of *concavin(−)* sporozoites showing clear invaginations of the IMC. **(B)** Exemplary SEM serial sections. Scale bar 500: nm

**Supplementary Figure S10.**
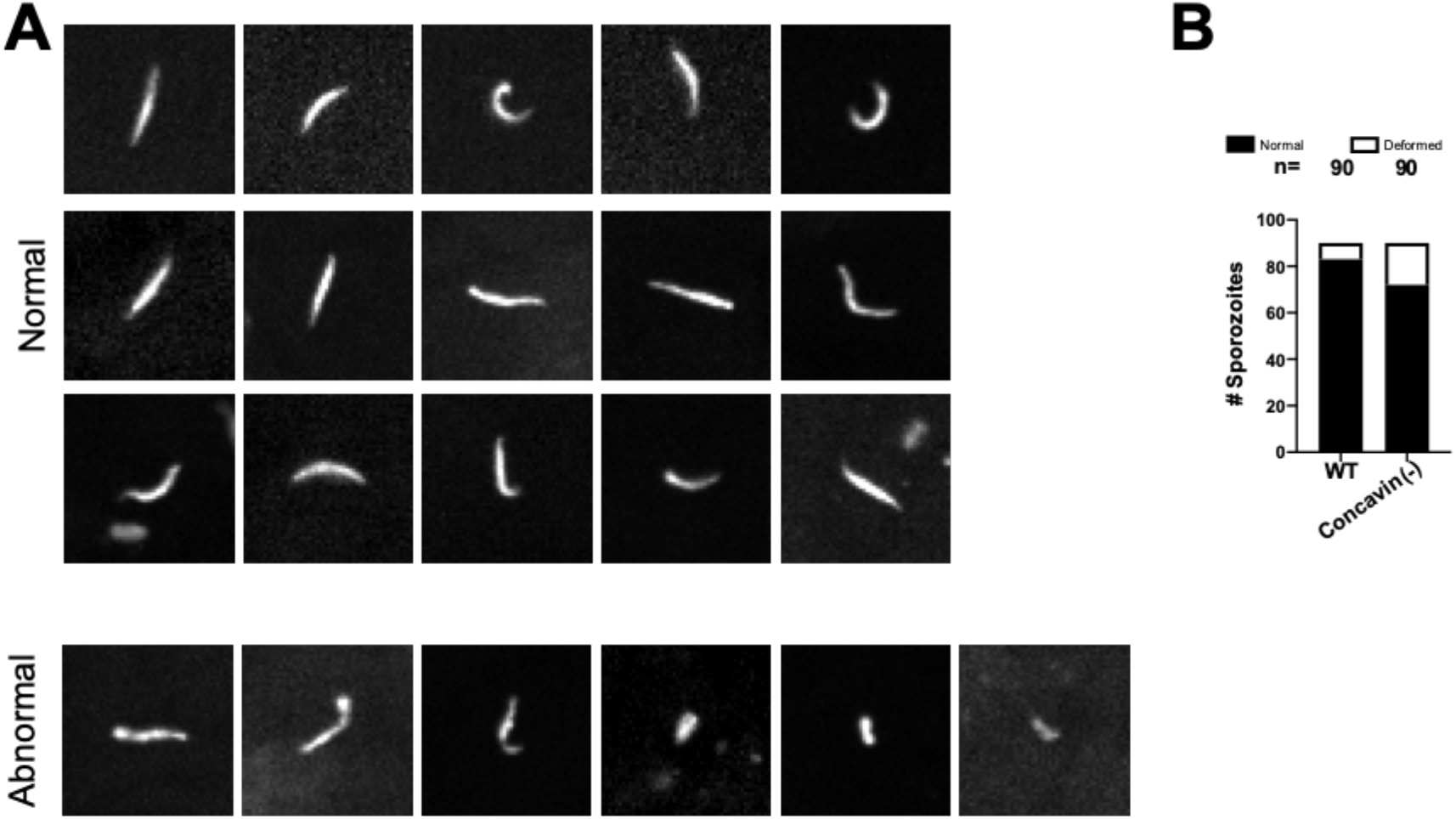
Examples from a representative bite site after transmission of *concavin(−)* parasites. **(A)** Morphology of normal and abnormal shaped concavin(−) sporozoites at the bite site. **(B)** Normally or abnormally shaped WT or *concavin(−)* sporozoites deposited in the skin. 90 sporozoites were observed for both parasite lines.

**Supplementary Table 1** Primer used for genotyping

**Supplementary Table 2** Raw data

**Supplementary Movie 1** FRAP of gliding concavin-GFP sporozoites

**Supplementary Movie 2** FRAP of gliding Phil1-GFP sporozoites

**Supplementary Movie 3** Sporozoite ejection on glass

**Supplementary Movie 4** Time-lapse of moving wild type and *concavin(−)* sporozoites in the skin

**Supplementary Movie 5** Collection of migrating *concavin(−)* sporozoites either deforming or disintegrating

