## Supplementary material for "Malaria transmission relies on concavin-mediated maintenance of *Plasmodium* sporozoite cell shape": Table S1

| Name | Sequence |
|---|---|
| JK 54 | AAAGCGGCCGCCGTTTTTCTTACTTATATATTTATACCAA |
| JK 55 | AAAGGTACCCGCATATCCTCATATATAATAAATTACCA |
| JK 56 | TTATCATAAAAGCTTGGCTGTCTT |
| JK 57 | AAAGCGGCCGCAACAAACAAATCTTCATGTTTGT |
| JK 58 | AAACCGCGGTGATATATGTACTCTTTTGTGTTCC |
| JK 111 | AACACCAGTCTGACACCAATTC |
| JK 112 | CCAGATCCAGTATTTTATACCATAGATG |
| JK 176 | GCAGCATTTTCTACTGGATAAGACAG |
| JK 177 | AAATTCGAAATGACAAACATTATAGAATGTACGTTCAAG |
| JK 178 | GGTTCCTTGTCCAATGGATATGACAAAG |
| JK 179 | AAATTCGAATTTGGAATATAACAAAAAAATATATCTCGTAATATA[illegible] |
| JK 236 | ATGACAAATGTTGTAGAAGCTACTTTTAAAACC |
| JK 237 | CTAGGATCCTTAGGCGCCTTTGTATAGTTCATCCATGCCATGT[illegible] |
| P 136 | CGCAATTTGTTGTACATAAAATAGGC |
| P 176 | CTAGACAGCCATCTCCATCTGG |
| P 177 | ATGCATAAACCGGTGTGTCTGG |
| P 210 | TTAACATCACCATCTAATTCAACAAG |
| P 322 | CCCCGTTGTCTGAGAAGG |
| P 587 | CTTTGGTGACAGATACTAC |
| P 668 | TGATTAGCATAGTTAAATAAAAAAAGTTG |
| P 862 | TCCAGTGAAAAGTTCTTCTCCT |
| P 961 | ATCCTCTGGTAATTTTTCG |
| P 963 | TAAGCAGTCGACCTACACAATCATGCATACTATGCC |
| P 969 | TAAGCAGAATTCCCCTTCCGAACAAATTTACGCC |
| P 970 | ATGGATCCaccaccaccaccaccaccaccaccCATATTATCTTTAGGGC |
| P 1446 | CAGCTGCTGGGATTACACATG |
